# Sleep-modulated disinhibition enables replay for memory consolidation, accelerated by ripples

**DOI:** 10.64898/2025.12.09.693276

**Authors:** Shrey Dutta

## Abstract

Memory consolidation involves elusive neural mechanisms. Here, we develop a biophysically detailed model of the entorhinal-hippocampal-cortical network to reveal that disinhibition drives synaptic and systems consolidation. Transitioning to slow-wave sleep via neuromodulatory dampening of inhibition generates up-down states and spontaneous, time-compressed replays of spatial sequences encoded with phase precession. Lateral inhibition levels unify physiological and pathological ripple diversity. Cortical disinhibition enables memory transfer from hippocampus. Weakened afferent CA1 synapses eliminate ripples but spare replays, proposing strategies to mitigate ripple disruptions. Replays sustain systems consolidation even without ripples, albeit slower; excessive weakening halts it, rescuable by enhanced hippocampal-to-cortical connectivity. Medial entorhinal cortex (MEC)-mediated CA1 disinhibition compensates for attenuated neuromodulatory CA1 disinhibition, mimicking MEC-input-dependent quiet wakefulness replay; under this configuration during wakefulness, artificially inducing disinhibition triggers replays and ripples that drive consolidation, underscoring disinhibition’s state-agnostic role. These insights elucidate disinhibition’s centrality in engraining memories and fostering hippocampal independence, reconciling empirical observations, yielding testable predictions, and identifying therapeutic avenues for memory disorders.

## Introduction

Memory consolidation, the process that transforms transient experiences into enduring memories, is fundamental to learning and cognition [1–5]. It encompasses synaptic consolidation—stabilizing memory engrams locally in the hippocampus [6–10]—and systems consolidation—integrating engrams with cortical networks [11–15]. Both rely on neural phenomena during slow-wave sleep [16–21] and quiet wakefulness [22–25]: memory replays reactivating sequential neural patterns from recent experiences [26–33]; sharp-wave ripples (SWRs), high-frequency oscillations that synchronize neuronal firing to promote synaptic plasticity [3, 34, 35]; and replay-mediated transfer redistributing hippocampal engrams to cortical regions (e.g., retrosplenial or prefrontal cortex), enabling long-term storage independent of the hippocampus [36–40]. Despite significant progress, the mechanisms initiating and regulating these phenomena remain elusive.

Emerging evidence reshapes our understanding of hippocampal SWR generation, focusing on disinhibition regulated by neuronal and neuromodulatory factors [41–43]. Traditionally, SWRs were thought to initiate sequential replay activity [44, 45]—but ripple-less replays [46] indicate independent mechanisms that remain to be elucidated. Moreover, although transient cortical disinhibition is necessary for hippocampal-to-cortical communication [47, 48], concurrent sleep-state alterations in inhibitory circuits [49–57] challenge its sufficiency, necessitating a refined mechanistic framework.

Studies indicate that SWRs facilitate hippocampal-to-cortical transfer of replayed sequences, as evidenced by co-occurrence with hippocampal replays [58–63] and ensuing cortical activity [64–67], along with impaired memory recall following ripple-associated replay disruption [68–73]. Wakeful ripple-associated replays tag engrams for selective sleep consolidation [74]. Ripple-less replay of non-salient trajectories [46] refines this view, since trajectory saliency dictates ripple coupling and thereby replay prioritization for cortical storage [75, 76]. These insights prompt pivotal questions: What mechanistic distinctions underlie ripple-associated versus ripple-less replay? Can ripple-less replay mediate hippocampal-to-cortical engram transfer?

To address these questions, existing models often omit biophysical integrations critical for multi-region dynamics across sleep-wake states. We thus develop a computational model of the entorhinal-hippocampal-cortical network, incorporating Hodgkin-Huxley neurons [77]; synaptic plasticity rules [78] including dynamic feedforward [79], feedback and lateral inhibition [80]; short-term plasticity [81]; inter- and intra-regional delays [82]; phase precession during encoding [83]; protein-mediated synaptic stabilization [84]; spine addition/removal [85]; and ionic dynamics for awake-sleep transitions [86]. This model simulates stimulus-triggered sequence encoding, spontaneous hippocampal replay and SWRs upon slow-wave sleep onset, and both synaptic and systems-level memory consolidation.

## Results

### Medial entorhinal cortex stimulation drives entorhinal-hippocampal spatial memory encoding

Our model includes neuronal populations for the medial entorhinal cortex (MEC), hippocampal subregions (dentate gyrus, DG; CA3; CA1), cortical regions (CR, e.g., prefrontal or retrosplenial cortex), and a background population that simulates asynchronous-irregular synaptic inputs during wakefulness [87, 88] and up-down dynamics during slow-wave sleep (SWS) [89] from unmodeled brain regions (Fig. 1a,b; Methods). To simulate the wakefulness-to-SWS transition, we lowered extracellular potassium buffer concentration ([K^+^]_Buffer_) from 3.5 to 3.0 mM, reflecting the observed drop in extracellular potassium concentration ([K^+^]_o_) during natural sleep [86]. This [K^+^]_o_ reduction decreases neuronal excitability via hyperpolarized resting potentials [86] and weakens inhibitory synaptic efficacy [49] via a K^+^-dependent disinhibition modulating factor (Methods), inducing SWS-typical up-down dynamics (Fig. 1b) [90].

**Fig 1:**
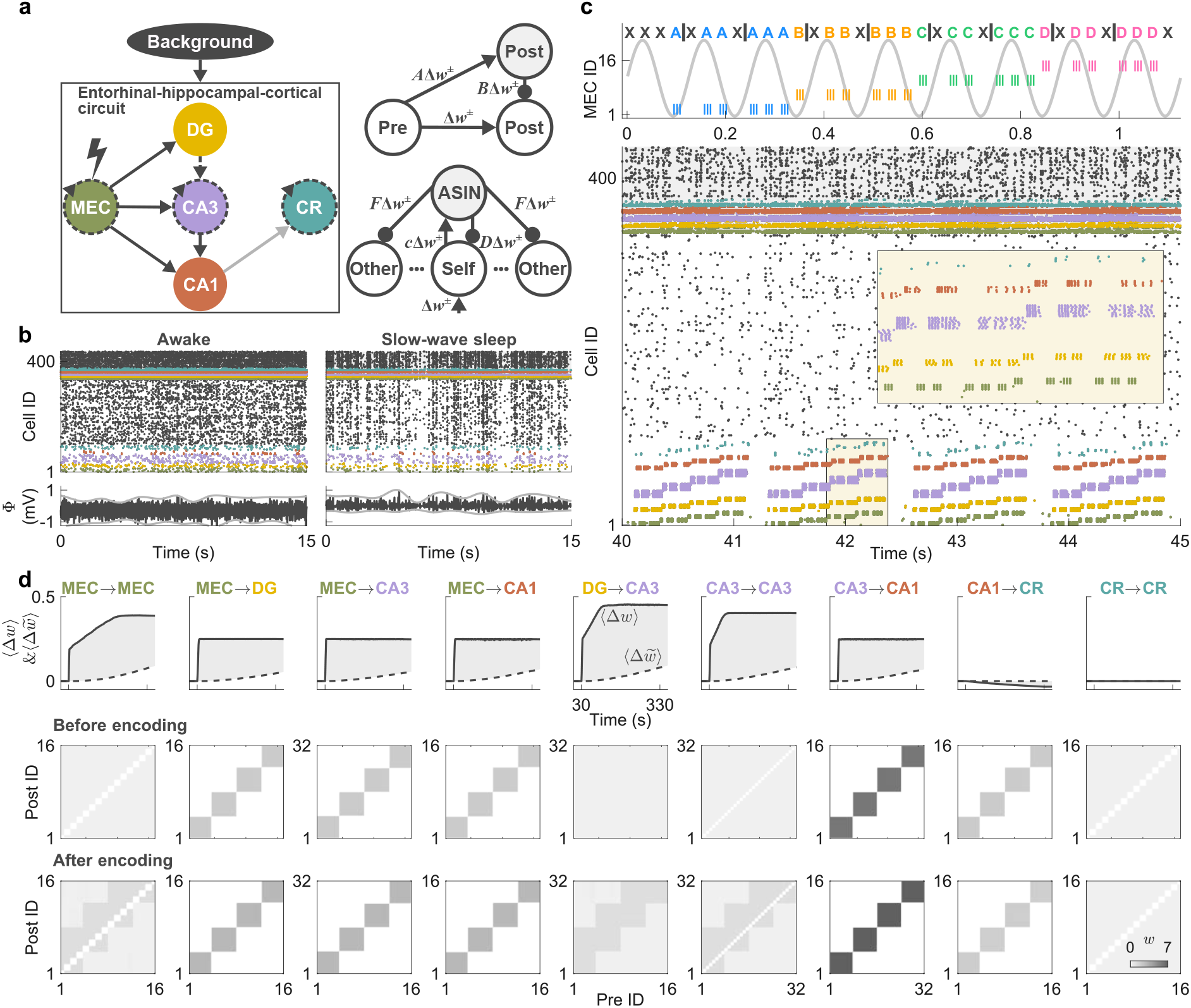
Entorhinal-hippocampal-cortical circuit model, awake/sleep dynamics, and stimulation-driven encoding. **a** Left: Schematic of the entorhinal-hippocampal-cortical circuit driven by background neural activity. Brain regions: medial entorhinal cortex (MEC, green), dentate gyrus (DG, yellow), CA3 (purple), CA1 (red), cortical regions (CR, blue-green), unmodeled brain regions (Background, black). Black solid arrows denote effective pre-encoding connections (MEC→DG, MEC→CA3, MEC→CA1, CA3→CA1). Gray solid arrow (CA1→CR) indicates anatomical connection with strong feedforward inhibition during wakefulness. Dashed arrows show ineffective (weak) pre-encoding connections (DG→CA3; recurrent in MEC, CA3, CR). Lightning bolt on MEC signifies stimulation initiating encoding. Right top: Dynamic synaptic updates for feedforward inhibition. Neurons: presynaptic excitatory (‘pre’, unfilled circle), postsynaptic excitatory (‘post’, unfilled circle) or inhibitory (gray filled circle). Updates: Δ*w*^*±*^ from ‘pre’ to excitatory ‘post’ drives *A*Δ*w*^*±*^ to ‘pre’→inhibitory ‘post’ and *B*Δ*w*^*±*^ to inhibitory ‘post’→excitatory ‘post’, where *A* = *a/M*, *B* = *b/M*, *M* is the number of interneurons in the feedforward inhibition path from ‘pre’ to ‘post’, and *a, b* are scaling constants. Right bottom: Dynamic synaptic updates for feedback and lateral inhibition. Neurons: excitatory in the assembly of ‘post’ (‘self’, middle unfilled circle), in other assemblies (‘other’, left/right unfilled circles), assembly-specific inhibitory neuron (ASIN, filled circle). Updates: Δ*w*^*±*^ to ‘post’ drives *c*Δ*w*^*±*^ to ‘post’→ASIN, *D*Δ*w*^*±*^ to ASIN→’self’ (feedback), *F* Δ*w*^*±*^ to ASIN→’other’ (lateral), where *D* = *d/N*, *F* = *f/N*, *N* is the number of neurons in the assembly, and *c, d, f* are scaling constants. Arrows: excitatory; round arrowheads: inhibitory. **b** Extracellular potassium buffer ([K^+^]_Buffer_) decrease (3.5 to 3.0 mM) transitions the dynamics from ‘awake’ to ‘slow-wave sleep’ via disinhibition. Top: Raster plots of spikes from 435 neurons (1–336 excitatory, 337–435 inhibitory) showing ‘awake’ (asynchronous-irregular; left) and ‘slow-wave sleep’ (up-down; right) dynamics before encoding. Bottom: Respective local field potential activity, (Φ), from ‘Background’ neurons. Gray lines trace the upper and lower envelopes. **c** Top: Stimulation pattern mimicking exploration from locations A to D with phase precession for 16 MEC neurons over 1.125 s. Each gray 8-Hz cycle (9 cycles, 125 ms each) is organized into four phases, with colored letters (blue A, orange B, green C, pink D: locations; black X: none) marking burst stimulation (three spikes) in the four neurons of each location-specific cell assembly, advancing to earlier phases with successive cycles. Bottom: Raster plot over 5 s during four complete stimulations of locations (MEC cell assemblies in green) ‘A’ to ‘D’. Inhibitory neurons at top with gray shading. Background activity (black) is asynchronous irregular (‘awake’). MEC stimulation drives downstream activity via effective connections (panel **a**). Inset: Zoomed view of the latter part of the second stimulation (boxed), showing delayed activation in downstream regions for locations C and D. **d** Top: Time course of changes (Δ) in mean excitatory-to-excitatory synaptic weights for connections in panel **a** during stimulation-driven encoding (⟨Δ*w*⟩, solid: fast overall [labile + stable] governed by spike-timing-dependent plasticity; 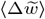, dashed: slow stable governed by protein-mediated stabilization; 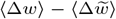, shading: labile component). The 5-min stimulation period (from 30 s to 330 s) represents repeated sequential exploration of locations ‘A’ to ‘D’. Middle & bottom: Matrices showing overall weights, *w*, between neurons before (middle) and after (bottom) the stimulation-driven encoding; white indicates no connection.

Excitatory-to-excitatory synaptic strengths evolve via triplet-spike-timing-dependent plasticity (triplet-STDP) [78], with proportional adjustments to associated inhibitory synapses for balanced feedforward, feedback, and lateral inhibition (Fig. 1a; Methods). Spatial navigation was simulated by sequentially stimulating location-specific MEC excitatory assemblies with phase-precessing spike bursts (Fig. 1c; Methods) [83]. Efferent MEC projections elicited delayed activation of corresponding assemblies in DG, CA3, and CA1, driving synaptic potentiation across regions (Fig. 1d) [91]. This strengthened initially weak MEC→MEC, DG→CA3, and CA3→CA3 synapses to encode sequence A–D, while enhancing MEC→DG/CA3/CA1 and CA3→CA1 projections. Strong CA1 CR feedforward inhibition [92–95] (Methods) limited CR activity below potentiation thresholds and thereby induced synaptic depression during exploration [96], preventing premature sequence maturation and allowing cortical engram recall only post-offline consolidation [38]. Potentiation saturates at peak spine receptor density but remains labile until protein-mediated synaptic stabilization (Methods).

Potentiated synapses stabilize over hours through lingering plasticity-related proteins (PRPs) from encoding, independent of engram reactivation [97]. To save stabilization time before sleep simulations, we assumed complete stabilization of encoding-induced changes by equating stable synaptic weights 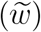 to overall (stable+labile) weights (*w*) and resetting PRPs to zero, enabling new spine formation and further potentiation during offline reactivation (Methods).

### Hippocampal disinhibition enables sequential replay of encoded engrams via inter-assembly delay

Following encoding, we transitioned to SWS dynamics. Reduced feedforward inhibition via a K^+^-dependent disinhibition factor (Methods) triggered spontaneous reactivation of location-specific assemblies in entorhinal and hippocampal regions (Fig. 2a). These reactivations replayed the encoded sequence (A–D) in compressed form, shortening from 1.125 s (Fig. 1c) to less than 400 ms (Fig. 2b–g), as observed in ethological contexts [33]. Replay initiated mainly during SWS up-phases, concordant with empirical studies [98]. Sequential replay propagated downstream [99–101], mirroring delayed hippocampal activation during encoding (Fig. 1c). Partial replays (length < 4) outnumbered full replays (length = 4) (Fig. 2h), recapitulating empirical patterns [102]. CA3 generates replay sequences independently of MEC inputs [101]; MEC silencing (via 60-fold feedforward inhibition increase) reduced replay occurrences but preserved lengths and durations, echoing empirical findings [101]. Intra-regional inter-assembly delays exceed intra-assembly delays (Methods); eliminating them (set to 0 ms) disrupted sequential replays (Fig. 2i), underscoring their role in sequence maintenance [103].

**Fig 2:**
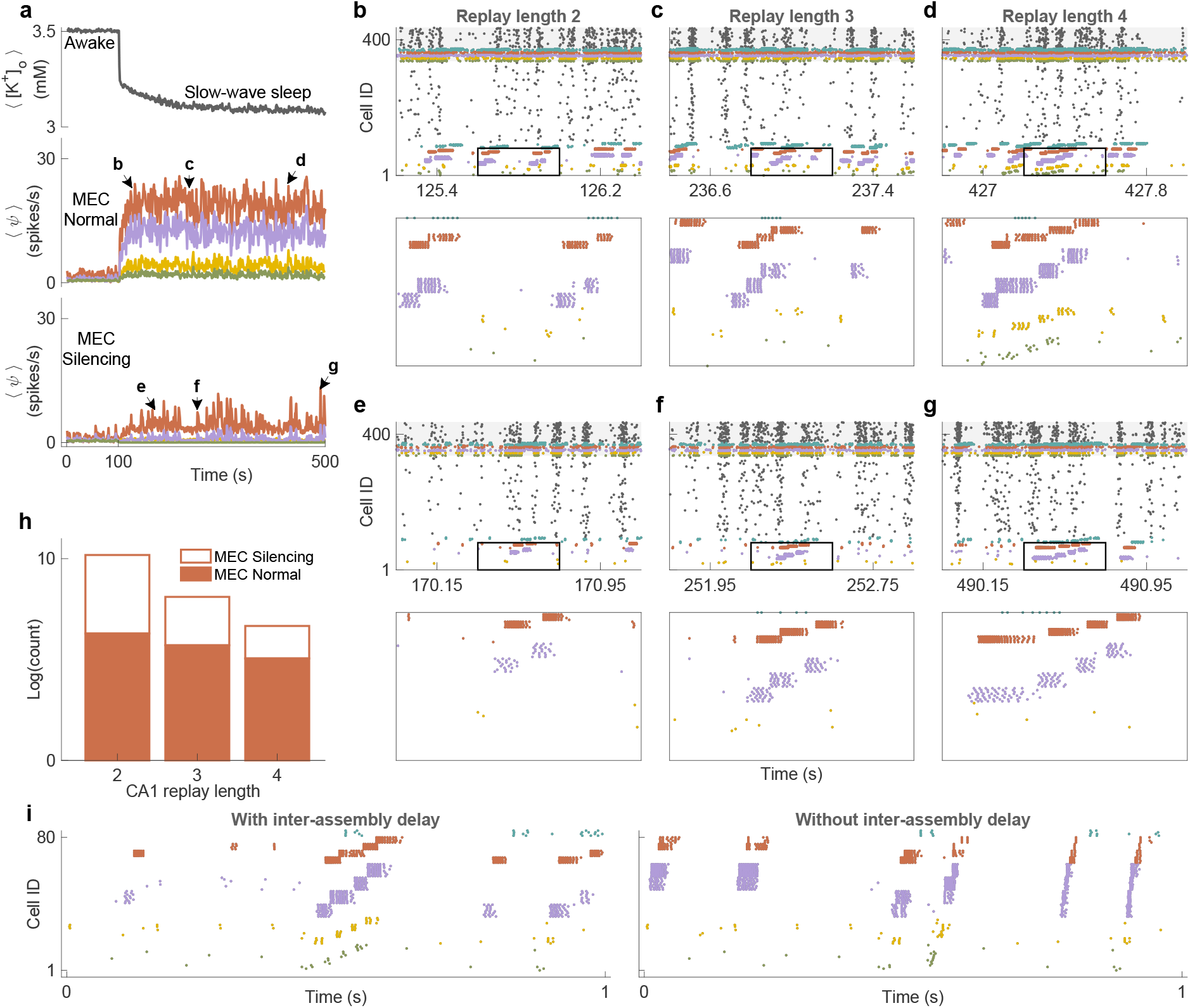
Transition to slow-wave sleep activates K^+^-dependent disinhibition that enables spontaneous time-compressed replay via inter-assembly delay. **a** Time courses of average extracellular K^+^,⟨[K^+^]o⟩, and average firing rate of entorhinal and hippocampal regions. Top: Temporal evolution of ⟨[K^+^]_o_⟩ from ‘Background’ neurons when [K^+^]_Buffer_ is decreased from 3.5 to 3.0 mM at 100 s to initiate the transition from ‘awake’ to ‘slow-wave sleep’ dynamics. Middle & bottom: Average firing rates, ⟨*ψ*⟩, during normal MEC dynamics (middle) and during silenced MEC dynamics by enhanced inhibition (bottom). Arrows labeled ‘b’ to ‘g’ point to the timestamps of rasters in panels **b** to **g. b–g** Top: 1200 ms raster plot of spikes from all neurons at the respective timestamps from panel **a**. Bottom: Zoom of the 400 ms box from the top panel showing exemplars of replays with lengths 2–4. **h** Bar plot showing the logarithm of total number of CA1 replays of lengths 2–4 during time-courses in panel **a. i** Left: Sequential replay with inter-assembly delay. Right: Disrupted replay sequence without inter-assembly delay. Colors represent brain regions as in Fig. 1.

### Reduced CA1 excitatory input yields ripple-less replay and decelerates synaptic consolidation, until disruption

Synchronized assembly bursts during replay evoke sharp-wave ripple (SWR)-like oscillations in simulated LFPs, originating in CA3 [45, 104–107] and propagating to CA1 [42, 104, 108, 109], as shown by frequency-time analyses (Fig. 3a). Motivated by ripple-less replays in non-salient contexts [46], we attenuated CA3→CA1 excitatory input without modeling saliency sources. This abolished ripples while preserving replay sequences (Fig. 3b), consistent with diminished ripples in salience-impaired states [110–114].

**Fig 3:**
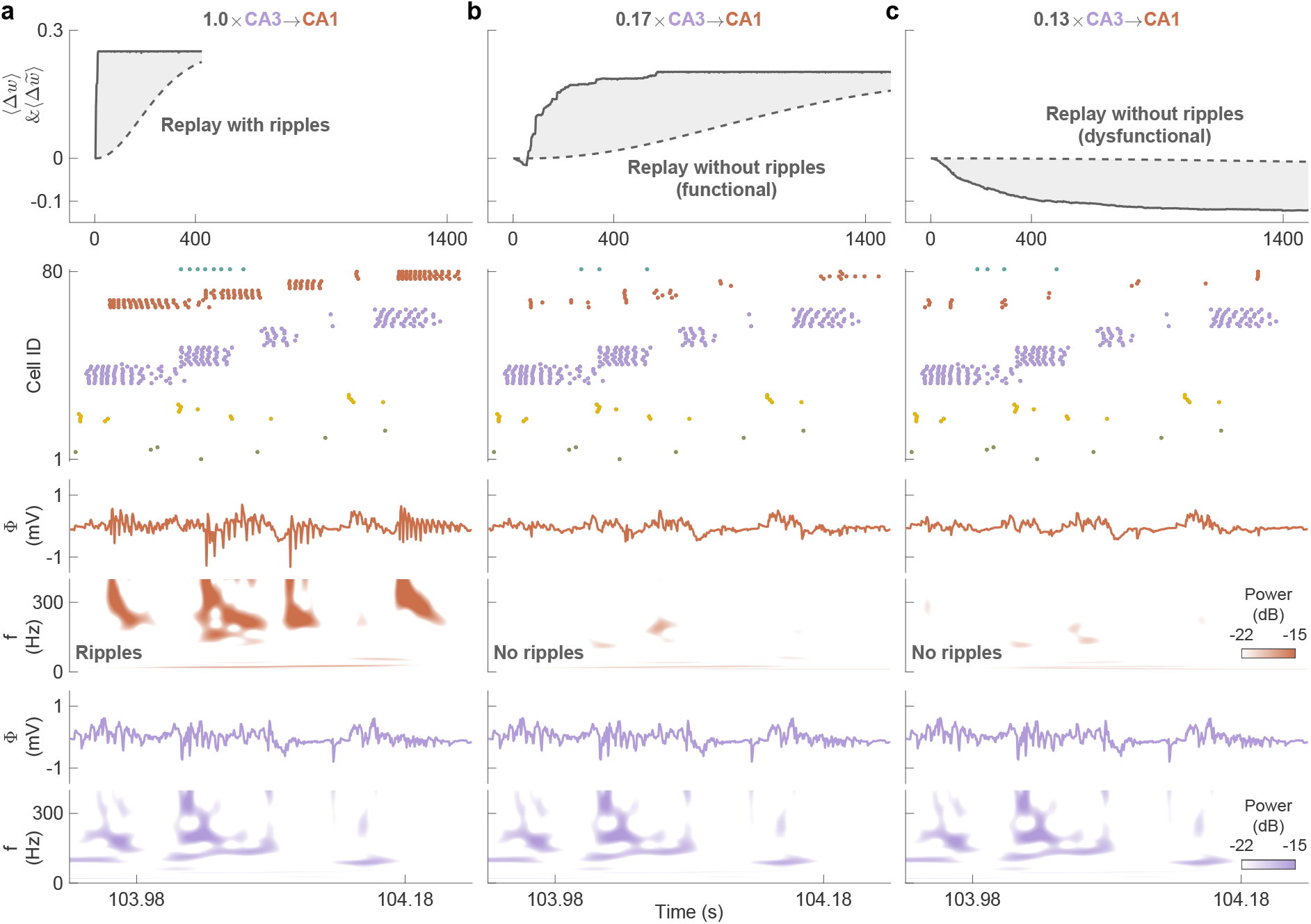
Reduced CA1 excitatory input renders replay without ripples, slowing synaptic consolidation, until disruption. **a–c** Dynamics with CA3→CA1 connectivity at 1.0× (a; with ripples), 0.17× (b; without ripples, functional), and 0.13× (c; without ripples, dysfunctional). Row 1: Time course of changes (Δ) in mean CA3→CA1 excitatory synaptic weights during slow-wave sleep (⟨Δ*w*⟩, solid, overall; 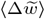, dashed, stable; shading, labile; see Fig. 1d for details). Rows 2–6: 300-ms raster plot (row 2), local field potential and time-frequency analysis for CA1 (rows 3–4) and CA3 (rows 5–6) showing an exemplar replay. Colors represent brain regions as in Fig. 1.

Although both ripple-associated and ripple-less replays consolidate hippocampal engram synapses by enhancing *w* via triplet-STDP and 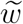 via PRP consumption (Fig. 3a,b; Methods), ripple-less replays at intermediate CA3→CA1 weight reduction (0.17×) consolidate slowly (functional), as triplet-STDP induces weaker potentiation at lower spike rates [78] and activity-dependent PRP synthesis (Methods) yields fewer PRPs for consumption (Fig. 3b). At higher reductions (0.13×), ripple-less replays persist but disrupt consolidation (dysfunctional; Fig. 3c).

### Lateral inhibition unifies empirically observed ripple diversity

To examine lateral inhibition (LI) effects at fixed reduced levels on CA1 ripple-associated replays, we disabled triplet-STDP and dynamic inhibitory updates to prevent interference with fixed LI values, while silencing MEC to isolate the CA3→CA1 circuit. LI modulation controls inter-assembly gap durations during sequence replay [115], with reductions that elevate average ripple power (Fig. 4a; Methods) [80]. At 100% LI, discrete ripple chains emerge as separable high-frequency bursts in frequency-time spectra, with successive events encoding adjacent assemblies (Fig. 4b, left), consistent with observed chains [99, 116, 117]. At 50% LI, shortened gaps concatenate fragments into prolonged ripples (Fig. 4b, middle), mirroring long-duration ripples [80, 105, 117]. At 0% LI, total fusion disrupts winner-takes-all dynamics [118–121], co-activating assemblies into disordered bursts (Fig. 4b, right), resembling ripples in schizophrenia models [122].

**Fig 4:**
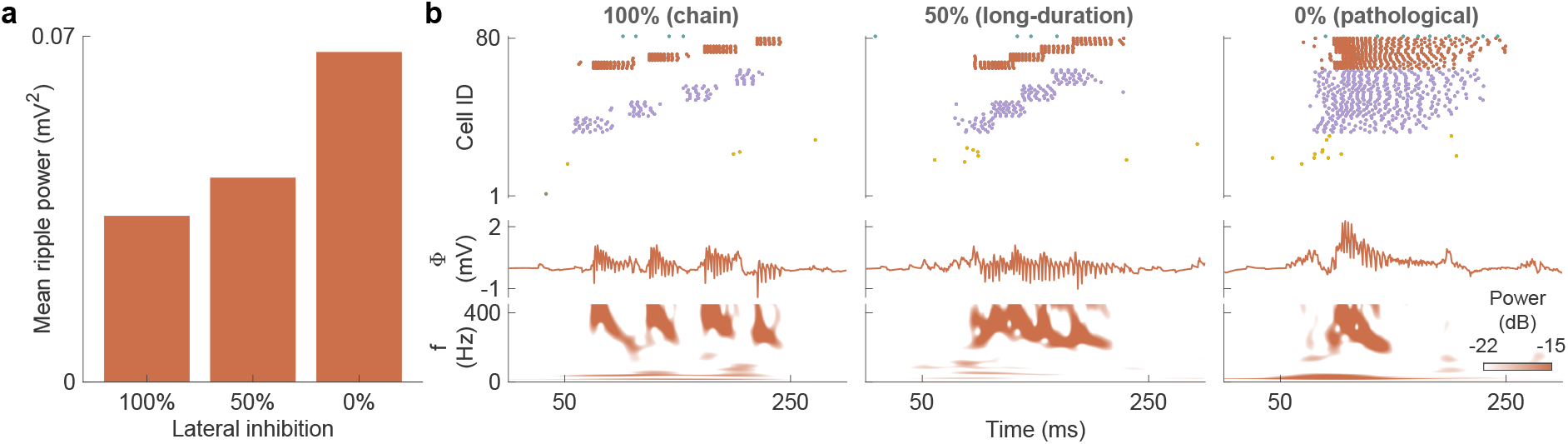
Different levels of lateral inhibition unify ripple diversity. **a** Mean power of CA1 ripple-associated replays (all lengths, including 1) increases as lateral inhibition (LI) decreases, computed over 200-s simulations. **b** 300-ms raster plot (row 1), CA1 local field potential (row 2), and CA1 frequency-time analysis (row 3) at 100% LI (ripple chain; left), 50% LI (long-duration ripple; middle), and 0% LI (pathological ripple; right). MEC is silenced to eliminate its inputs to CA3 and CA1, thereby isolating CA3 modulation of CA1. Colors represent brain regions as in Fig. 1.

### Cortical disinhibition enables sequence transfer from hippocampus resulting in hippocampal independence

Systems consolidation involves the transfer of hippocampal memories to cortical regions (CRs) [39, 123–126], rendering them independent of the hippocampus [11, 38, 127, 128] for endurance despite hippocampal forgetting [13, 38, 129–131]. Sleep-modulated CR disinhibition enables CA1 ripple-associated replay to overcome strong feedforward inhibition active during wakefulness, which sequentially activates CR cell assemblies, thereby potentiating synapses to encode the transferred sequence (Fig. 5a) [55,57,132]. CR displays spontaneous replays even after silencing entorhinal and hippocampal regions, exhibiting hippocampal independence and confirming successful transfer.

**Fig 5:**
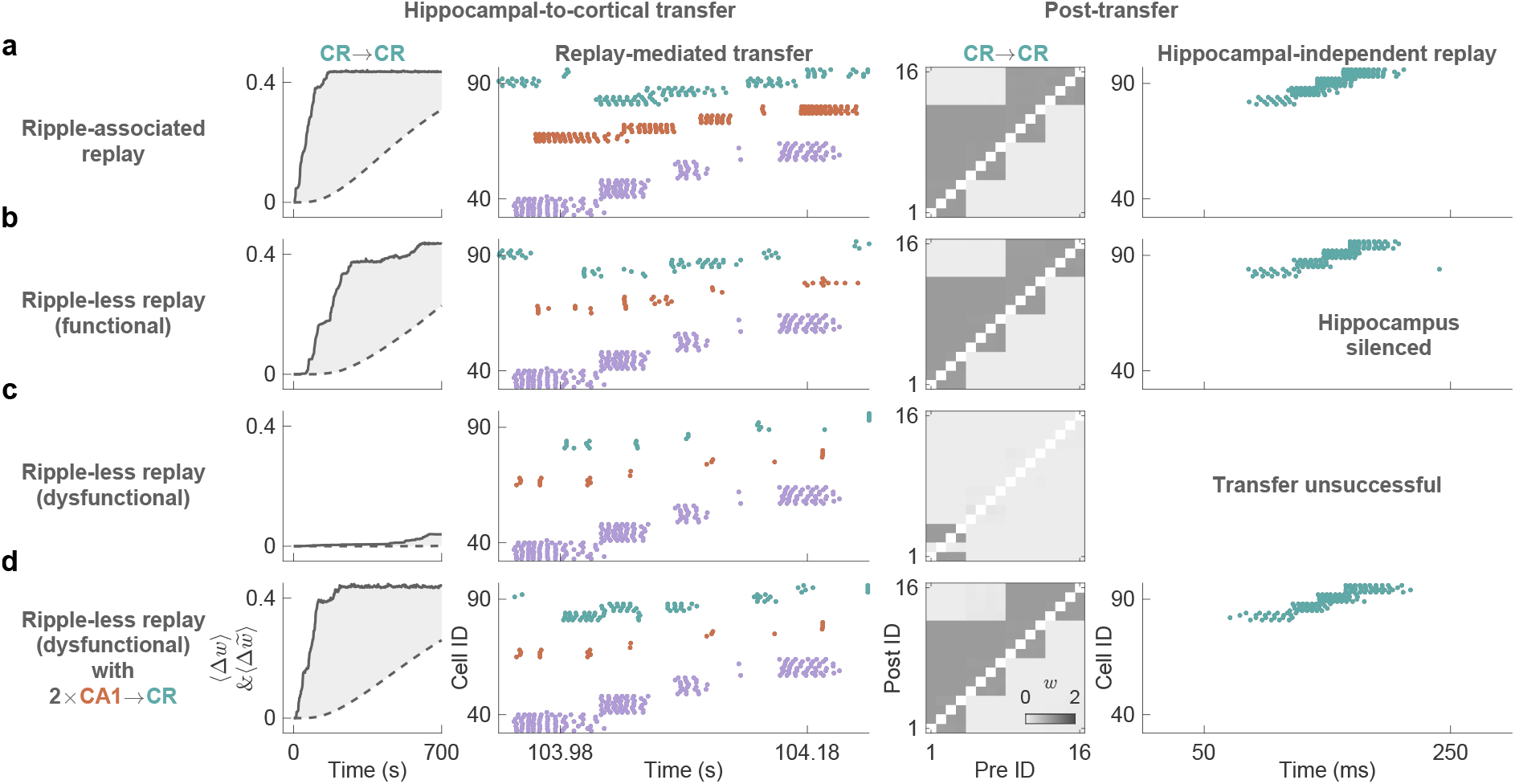
Hippocampal-to-cortical transfer via ripple-associated and ripple-less replay enables hippocampal independence. **a–d** Dynamics under ripple-associated replay (panel **a**), functional ripple-less replay (panel **b**), dysfunctional ripple-less replay (panel **c**), and dysfunctional ripple-less replay with doubled CA1→CR excitatory strength (panel **d**). Columns 1–2 trace the transfer: time course of changes (Δ) in mean CR→CR excitatory weights (column 1); 300-ms raster plot from Fig. 3 including CR neurons, depicting replay-mediated memory transfer to cortical assemblies (column 2). Columns 3–4 reveal post-transfer cortical autonomy: CR→CR excitatory weight matrix encoding transferred engram (column 3); hippocampal-independent replay after silencing MEC, DG, CA3, and CA1 through 60-fold feedforward inhibition increase, underscoring self-sustained CR replay (column 4). Colors denote brain regions as in Fig. 1.

Functional CA1 ripple-less replay transfers successfully to CR but more slowly than ripple-associated replay (Fig. 5b). However, dysfunctional ripple-less replay fails to achieve systems consolidation (Fig. 5c), but doubling CA1→CR excitatory weights restores consolidation (Fig. 5d), with implications for salience-impaired conditions where disrupted ripples hinder this process [110–114, 133].

### Absent CA1 disinhibition explains MEC-input-dependent quiet wakefulness replay via MEC-mediated disinhibition

To explain why extended quiet wakefulness (QW) replay in hippocampal CA1 requires direct MEC inputs [99], we ablated MEC→CA1 excitatory and inhibitory connections. Despite this, CA3-initiated replay propagated to CA1, recapitulating MEC-independent sleep replay [99] (Fig. 6a).

**Fig 6:**
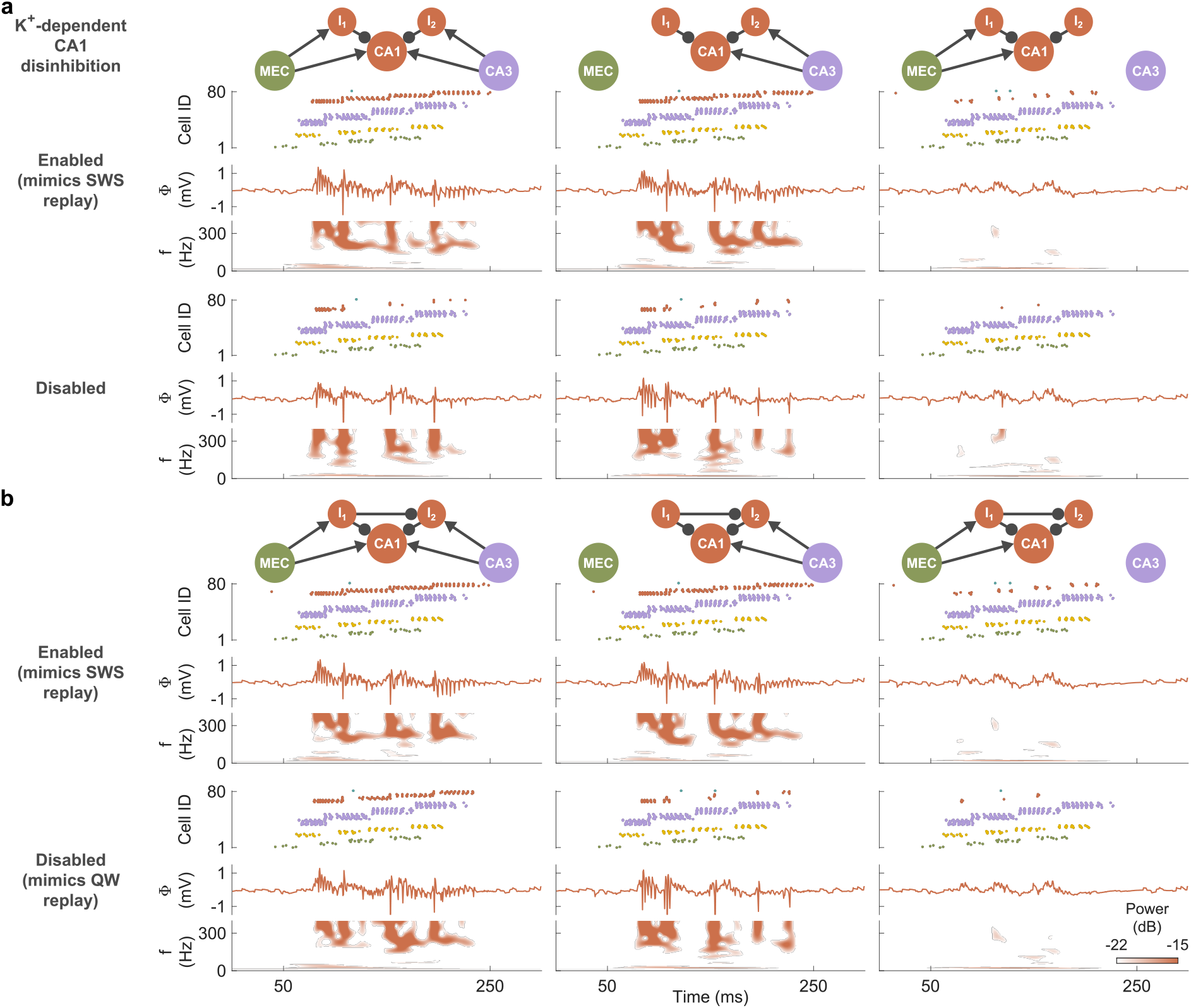
Disabling K^+^-dependent CA1 disinhibition underlies MEC-input-dependent quiet wakefulness replay through MEC-mediated disinhibition. **a**,**b** Row 1: Schematics of feedforward excitatory inputs (arrows) from MEC and CA3 (column 1), CA3-only (column 2), or MEC-only (column 3) to CA1 excitatory and inhibitory neurons; absent (panel **a**) or present (panel **b**) inhibitory link (round arrowheads) from MEC-to CA3-receiving CA1 interneurons (I_1_→I_2_ = 50), enabling MEC-mediated disinhibition of excitatory CA1 neurons and gating CA3 input to CA1. Rows 2–7: Replay dynamics, with K^+^-dependent disinhibition enabled (rows 2–4) or disabled (rows 5–7), in 300-ms raster plots (rows 2, 5), CA1 local field potentials (rows 3, 6), and time– frequency analyses (rows 4, 7). Colors denote brain regions as in Fig. 1.

In QW, neuromodulation-associated disinhibition is lower relative to sleep [57, 134, 135]. Disabling CA1 disinhibition eliminates replay propagation, irrespective of MEC and CA3 inputs (Fig. 6a). This conflicts with empirical observations where combined MEC and CA3 inputs enable QW replay, prompting the inclusion of MEC-mediated disinhibition to compensate for the absent sleep-modulated disinhibition. We connected MEC-recipient CA1 interneuron to inhibit CA3-recipient interneuron, inducing transient feedforward disinhibition (Fig. 6b) [136–138]. This restores CA3-elicited CA1 replay under disabled CA1 disinhibition but with concurrent MEC inputs, thus replicating MEC-input dependence in QW replay dynamics (Fig. 6b). Restoring sleep-modulated CA1 disinhibition eliminates MEC-input requirement while retaining CA3 dependence, matching empirical sleep replay [99].

### Artificially induced disinhibition during wakefulness triggers memory processes driving consolidation

Memory replay, SWRs, and hippocampal-to-cortical transfer, typically sleep-associated, also occur during QW [123, 139–142]. To assess if disinhibition suffices to drive these memory processes independent of sleep-specific dynamics, we artificially induced disinhibition in entorhinal, hippocampal, and cortical regions under the derived QW replay parameters—disabled CA1 disinhibition and enabled MEC-mediated CA1 disinhibition— while preserving wakeful asynchronous background activity. This triggers spontaneous replays that achieve synaptic consolidation and hippocampal-to-cortical transfer for systems consolidation (Fig. 7), recapitulating sleep-associated processes. Thus, sustained disinhibition, not SWS up-down states [98], is fundamental across brain states for eliciting replay, ripples, and hippocampal-to-cortical transfer [55, 57, 132].

**Fig 7:**
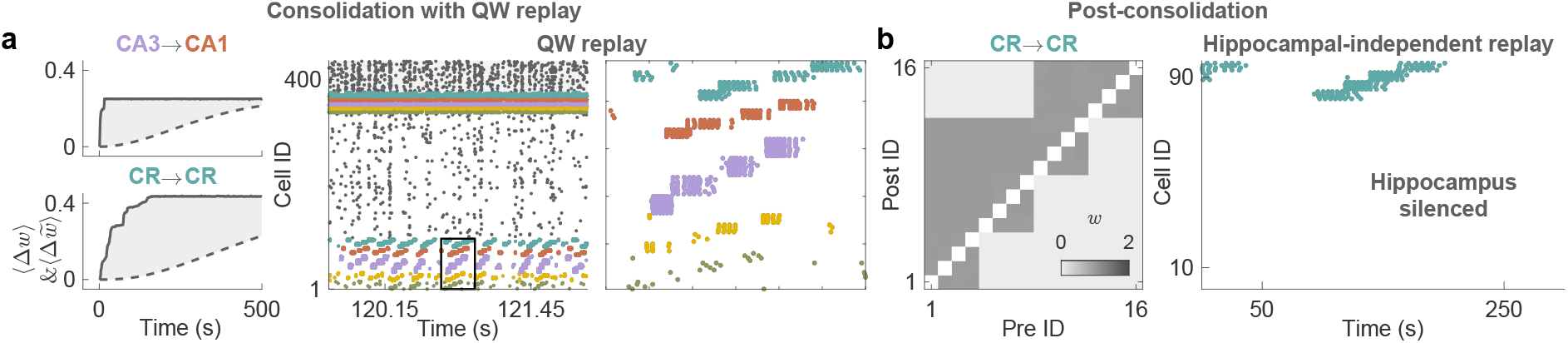
Artificially induced disinhibition under derived QW replay parameters drives synaptic and systems consolidation, yielding hippocampal independence. **a** QW replay-mediated consolidation: time course of changes (Δ) in mean synaptic weights of CA3→CA1 (top left) and CR→CR (bottom left); 2300 ms raster plot of spikes from all neurons (middle); zoom of the 300 ms box from the middle panel (right). **b** Post-consolidation cortical autonomy: CR→CR synaptic weight matrix encoding transferred engram (left); hippocampal-independent CR replay upon silencing MEC, DG, CA3, and CA1 (right). Colors denote brain regions as in Fig. 1.

## Discussion

Our computational model reveals how sleep-modulated disinhibition coordinates memory consolidation through spontaneous replay events that promote synaptic strengthening and systems-level transfer. We mimic slow-wave sleep (SWS) by reducing extracellular potassium concentration ([K^+^]_o_), which lowers neuronal excitability [86] via a shift from high-to low-conductance states [143, 144]. This reduction diminishes neuromodulatory tone [145], thereby weakening inhibition [50,56] and initiating up-down states [90]. Instead of disfacilitation [146] or inhibitory buildup [147] to terminate up states, with resurgence via recovery from disfacilitation [148] or rebound bursts [149], our model generates up states via disinhibition and terminates them through short-term depression [150] under baseline low-[K^+^]_o_ conditions. This posits that up-down states emerge from homeostatic compensation for sleep’s low excitability and firing-rate reductions via disinhibition [151]— an activity-dependent neuromodulatory process that may presage persistent maladaptive disinhibition in disorders such as Alzheimer’s disease [152], epilepsy [153], and others [154–156], wherein chronic low activity triggers inhibitory weakening via synaptic remodeling. This homeostatic view, echoed in computational studies [157–160], proposes in vivo experiments inducing low excitability—via pharmacological Na^+^ channel blockade [161], genetic overexpression of Kir4.1 in astrocytes [162], or optogenetic neuronal hyperpolarization [163]—to verify if this elicits disinhibition [164] and up-down states independently of natural sleep. In Alzheimer’s disease, astrocyte dysfunction disrupts K^+^ buffering [165], correlating with SWS fragmentation that accelerates pathological progression [166]; our K^+^ buffering model, supported by optogenetic astrocyte interventions restoring sleep rhythms [167], suggests that boosting astrocytic Kir4.1 function may stabilize SWS cycles and ameliorate memory deficits, given that its impairment aggravates disease features [168].

Complementing prior models of slow synaptic weight stabilization [158, 160], our offline consolidation integrates activity-dependent protein synthesis for slow-timescale engram stability [84] and spontaneous replays that engage fast-timescale triplet-STDP for selective engram strengthening [169]. The model’s sustained disinhibition—distinct from transient variants in ripple initiation [42, 43, 170] and gating hippocampal-to-cortical input [48]—engages dedicated pathways for engram sequence replay, sharp-wave ripple (SWR) generation, and hippocampal-to-cortical transfer. In contrast to views positing SWRs as replay initiators [44, 45], we identify independent mechanisms: inter-assembly delays drive sequential reactivation, while lateral inhibition preserves assembly separation, enabling robust replay despite ripple absence (ripple-less replays [46]) upon CA1 excitatory afferent attenuation. Synaptic and systems consolidation persist in ripple-less replay, albeit at reduced rates, motivating empirical assays of selective ripple ablation via CA1 afferent weakening—unlike prior methods co-disrupting replays and ripples [68–73]—to disentangle their roles. Paralleling ripple-less replays of non-salient trajectories [46], these mechanisms suggest tunable adjustment of CA1 afferent strength to normalize ripple dynamics in salience-impaired conditions: enhancing where ripples diminish, as in Alzheimer’s disease [110,111], epilepsy (physiological SWRs) [112,113], and aging [110,114]; attenuating where they surge, as in schizophrenia [122, 133, 171]. Amid hippocampal seizure risks [172], alternatively strengthening CA1-to-cortical connectivity restores engram transfer even with dysfunctional ripple-less replay. Future extensions could incorporate saliency cues from locus coeruleus (novelty) [173], ventral tegmental area (reward) [174], basal amygdala (emotion) [175], or lateral entorhinal cortex (contextual cues stabilizing ensembles) [176–178] to prioritize encoding of internally imagined [179] or externally sensed [174] events that deviate from anticipated offline replay [180, 181].

LI’s modulation of inter-assembly gap duration during simulated sequence replay underlies ripple diversity: strong LI yields discrete chains (doublets, triplets) that link sequence fragments [99, 116, 117]; intermediate LI concatenates them into prolonged events preserving replay sequence [80, 105, 117]; weak LI induces fusion and sequence erosion—mirroring order-ablated patterns from parvalbumin interneuron silencing [40] and pathological disinhibition in schizophrenia [122,182]—positioning LI as a prime therapeutic target for restoring trajectory fidelity.

Neuromodulatory changes similar to those in SWS occur during quiet wakefulness (QW), albeit to a lesser extent [134], reducing inhibitory tone [57, 135]. This disinhibition potentially explains delayed replays of recent experiences [31] by overcoming learning-induced inhibition. Our model reveals that MEC-mediated disinhibition compensates the disabled CA1 disinhibition to gate CA3 input and trigger ripple-associated replay, mirroring empirical MEC-input dependence during QW [31, 99]; whereas enabling CA1 disinhibition eliminates this MEC-input dependence, as seen in SWS replay [99]. This suggests reduced neuromodulator-driven CA1 disinhibition during QW relative to SWS, testable by optogenetically enhancing CA1 disinhibition to determine if MEC-input dependence is eliminated, akin to sleep dynamics [99]. Future models could incorporate CA1 recurrent connections [183] to explain MEC- and CA3-independent CA1 assembly sequences during active exploration [184], potentially triggered by contextual cues from lateral entorhinal cortex [176] and thalamic nucleus reuniens [185]. While SWS up-states create excitatory windows facilitating hippocampal ripples [186], our model demonstrates that artificially induced sustained disinhibition during wakefulness—independent of up-states—suffices to initiate and propagate ripple-associated replay to cortical regions [48, 141], providing a mechanistic basis for the homeostatic trade-off of hippocampal SWR events between quiet wakefulness and SWS [187] via disinhibition. This implies nuanced roles for cortical SWS up-states and associated sleep spindles, such as integrating transferred information across broader cortical networks [47], wherein spindle absence—by disrupting hippocampal input gating and inter-cortical integration—may drive consolidation deficits under sleep deprivation [188], a mechanism ripe for future modeling.

Although our model captures key hippocampal replay dynamics through recurrent MEC and CA3 connections that encode sequential engram and initiate offline replay [61,189,190], it omits feedback projections, including cortical input for replay selection [39, 191] and CA3→DG connectivity for dentate sharp wave [192]. Instead, feedforward input from MEC replay [189] yields underexplored DG patterns that recapitulate encoding-like sequential activation [193, 194]. Extending MEC phase precession could model reverse replay, with past assembly following current one to form backward association [195], abstractly resembling symmetrical STDP [61] despite asymmetric rules. The model couples inhibitory plasticity updates to excitatory-to-excitatory plasticity for excitation-inhibition balance, yet biologically observed independent local inhibitory rules [196] may intrinsically maintain this balance in future versions. During encoding, CA1 activity evokes transient cortical activation incapable of long-term potentiation [96, 197], modeled via heightened feedforward inhibition, preventing premature cortical engram maturation [38]; refining short-term synaptic plasticity to strengthen cortical synapses on working memory timescales [198, 199] could elucidate working memory’s role in systems consolidation [200]. Broader extensions could explore preplay for engram recruitment [201] probably shaped by past experience [202]; rapid eye movement engram dynamics for complementary consolidation [50]; larger networks for multi-memory storage [88, 158]; mechanisms underlying replay speed increases for longer trajectories without extending duration [33]; and gradual forgetting [13, 38, 203] via neurogenesis-driven processes [129, 204–206] that facilitate systems reconsolidation to update memories [205], increasing selectivity [160], and promoting generalization with low-dimensional concept retention [12, 15, 207–210].

## Methods

### Individual neuronal dynamics

The core dynamics of each neuron revolve around its membrane potential *V* (in mV), the electrical voltage across the cell’s membrane that governs action potential initiation, described by the stochastic Hodgkin-Huxley equation [77, 211]:

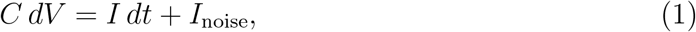

where *C* (in *µ*F/cm^2^) is the membrane capacitance, *dV* is the infinitesimal change in *V*, *dt* is the infinitesimal time interval, *I* = −*I*_Na_ −*I*_K_ −*I*_Cl_ −*I*_syn_ is the deterministic current (in *µ*A/cm^2^) comprising the sodium current *I*_Na_ (usually inward, initiating depolarization), the potassium current *I*_K_ (usually outward, enabling repolarization), the chloride leak current *I*_Cl_ (usually outward at rest, stabilizing baseline *V*), and the synaptic current *I*_syn_ (inward for excitatory or outward for inhibitory signals), and *I*_noise_ = *σ*_noise_*C dW* is the stochastic current with *dW* (in 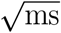) normally distributed with mean zero and variance *dt*, and *σ*_noise_ (in 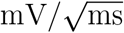) scaling the noise amplitude. This stochastic term approximates net current perturbations from thermal fluctuations driving random ion channel gating [212] and synaptic neurotransmitter release [213]. The equation captures how depolarizing/hyperpolarizing current balance, augmented by noise, drives *V* dynamics and action potential firing when depolarization prevails.

The ionic currents flow through specialized protein channels in the membrane, modeled as

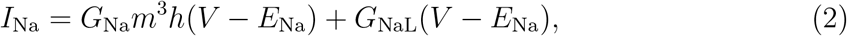

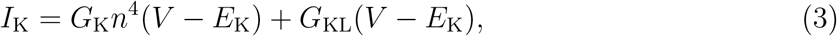

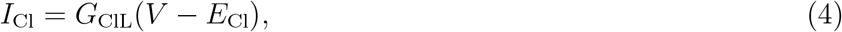

where *G*_Na_ (in mS/cm^2^) and *G*_K_ are the maximum possible flow rates (conductances) through sodium and potassium channels when they are fully open during an action potential. *G*_NaL_, *G*_KL_, and *G*_ClL_ are small, constant leak conductances that allow baseline ion flow, contributing to the neuron’s resting membrane potential. The variables *m, h*, and *n* are fractions between 0 and 1 that represent the proportion of channels in an open or active state: *m* increases to open sodium channels as *V* becomes less negative (depolarizes), *h* decreases to inactivate sodium channels and limit the duration of the action potential, and *n* increases with a delay to open potassium channels and return *V* to rest. The exponents (such as *m*^3^*h*) reflect that multiple molecular subunits must align for a channel to open fully. *E*_Na_, *E*_K_, and *E*_Cl_ are the reversal potentials for each ion, the specific voltage values at which the net flow of that ion stops because electrical and concentration forces balance (typically positive for *E*_Na_ favoring inward flow, and negative for *E*_K_ and *E*_Cl_ favoring outward flow).

The fractions *m, h*, and *n* adjust dynamically over time based on *V* :

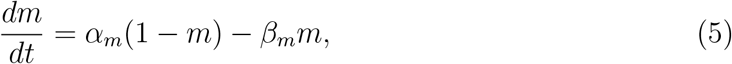

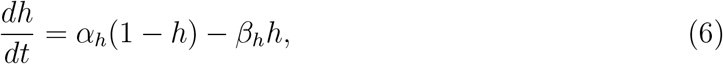

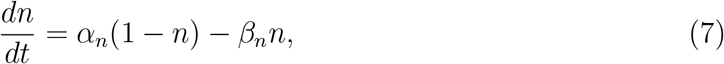

where these equations describe how the fractions approach steady states: the *α* terms increase the fraction toward 1 (opening or activating the channel), while the *β* terms decrease it toward 0 (closing or inactivating the channel). This allows channels to open quickly during depolarization and close during repolarization. The rates *α* and *β* vary with the membrane potential *V* [214, 215]:

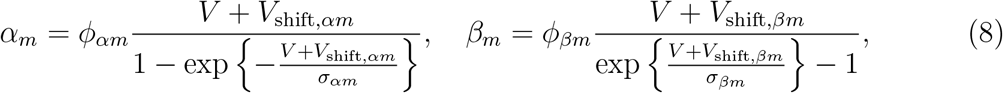

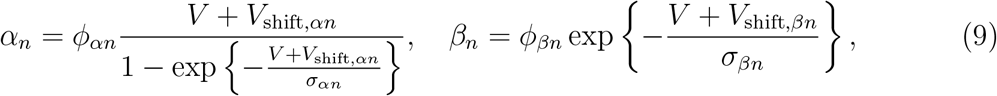

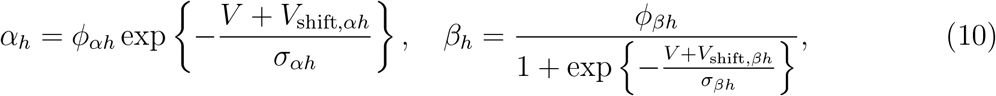

where the *ϕ* parameters (in ms^−1^) set the baseline magnitude of each rate, *V*_shift_ adjusts the voltage level at which the rate reaches half its maximum to align with typical neuronal firing thresholds (e.g., −50 mV), and *σ* determines how sensitively the rate responds to small changes in *V* (smaller values mean sharper responses).

The reversal potentials depend on ion concentrations across the membrane, following the Nernst equation:

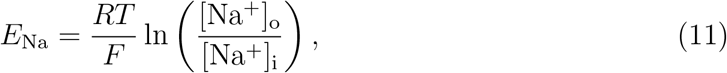

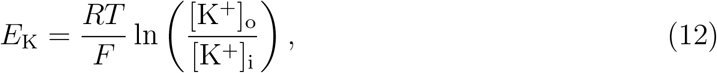

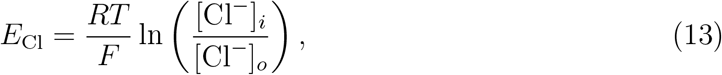

where *R* (in J/mol·K) is the universal gas constant, *T* (in K) is the absolute temperature, and *F* (in C/mol) is Faraday’s constant. Chloride concentrations are held constant for simplicity, while sodium and potassium levels vary with neuronal activity, as described below.

Ion concentrations evolve as ions move through channels during action potentials and are restored by pumps [214]:

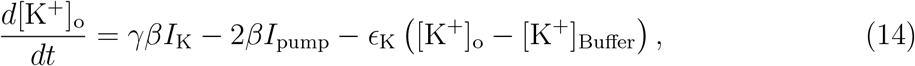

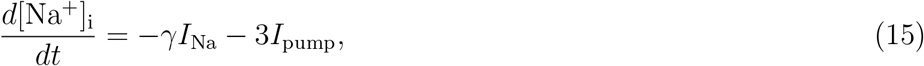

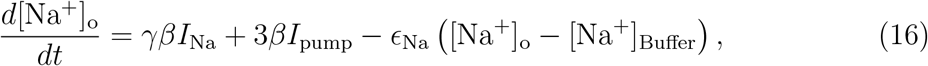

where *γ* is a conversion factor that translates electrical current measurements (in *µ*A/cm^2^) into changes in ion concentration (in mM/s), based on cell size and charge per ion; *β* is the ratio of the cell’s internal volume to the surrounding external space, reflecting that the external space is smaller and thus more prone to rapid concentration shifts; and *I*_pump_ is the current generated by the sodium-potassium pump, a protein that actively moves 3 sodium ions out and 2 potassium ions in per cycle to counteract imbalances from firing. The terms with *ϵ*_K_ (in s^−1^) (potassium clearance rate) and *ϵ*_Na_ (sodium clearance rate) model diffusion and buffering that pull external potassium and sodium toward [K^+^]_Buffer_ and [Na^+^]_Buffer_, respectively—stable reference concentrations representing the combined stabilizing effects of nearby blood vessels, tissues, and support cells like astrocytes.

Assuming total ion conservation [214, 215], achieved by balancing inward Na^+^ flux with outward K^+^ flux, the intracellular K^+^ concentration, [K^+^]_i_, is given by:

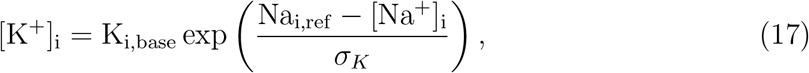

where K_i,base_ is the resting intracellular K^+^ level; Na_i,ref_ is a reference intracellular Na^+^ level; and *σ*_*K*_ is the exponential scale factor that ensures [K^+^]_i_ remains positive as [Na^+^]_i_ increases.

The pump current, *I* _pump_, increases gradually (in a sigmoidal pattern) as [Na^+^]_i_or [K^+^]_o_levels rise, mimicking how the pump ramps up to restore balance:

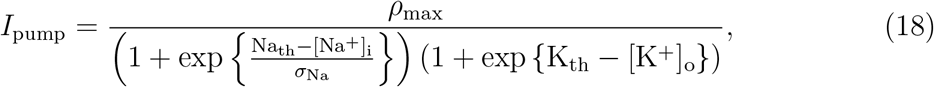

where *ρ*_max_ (in mM/s) is the pump’s maximum operating speed, Na_th_ and K_th_ are the concentration levels at which the pump reaches half-speed, and *σ*_Na_ controls how abruptly the pump responds to sodium increases.

The synaptic current, *I*_syn_, aggregates inputs from *P*^ex^ presynaptic excitatory neurons [216–219]:

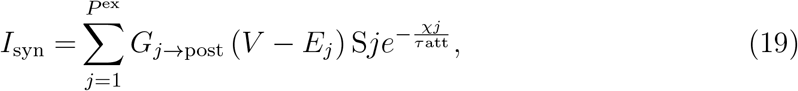

where *G*_*j*→post_ is the maximum synaptic conductance for synapses between postsynaptic (excitatory or inhibitory) and the *j*^th^ presynaptic (excitatory or inhibitory) neurons, *E*_*j*_ is the reversal potential (typically positive for excitatory, producing inward depolarizing currents at resting potential; negative for inhibitory, producing outward hyperpolarizing currents at resting potential), *S*_*j*_ is the fraction (ranging from 0 to 1) of open postsynaptic channels at the synapses formed with the *j*^th^ presynaptic neuron, and the exponential term attenuates synaptic strength when the presynaptic neuron is in depolarization block, with *χ*_*j*_ integrating a voltage-dependent measure of the duration of the depolarization block and *τ*_att_ setting the attenuation scale. *S*_*j*_ depends on the presynaptic neuron’s voltage, *V*_*j*_, following first-order kinetics [215–217]:

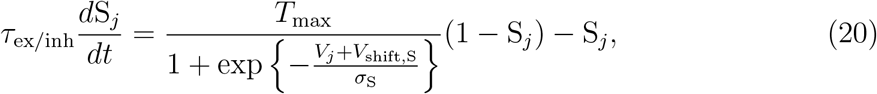

where *τ*_ex*/*inh_ is the decay time scale for excitatory/inhibitory synapses, *T*_max_ is the maximum neurotransmitter release rate, *V*_shift,S_ adjusts the presynaptic voltage for half-maximal release, and *σ*_S_ sets how sharply release responds to *V*_*j*_ changes. The dynamics of *χ*_*j*_ are governed by:

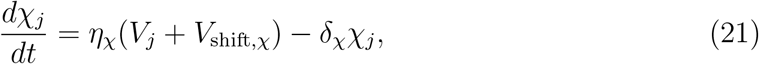

where *η*_*χ*_ equals *η*_max_ if the presynaptic membrane potential *V*_*j*_ falls within the depolarization block range (*V*_dep,low_ to *V*_dep,high_) and 0 otherwise, *V*_shift,*χ*_ is a voltage shift that modulates the buildup rate, and *δ*_*χ*_ is the decay rate of *χ*_*j*_ that restores synaptic strength post-block and prevents overload during prolonged block.

Maximal synaptic conductances, *G*_pre→post_, depend on short-term and long-term plasticity states:

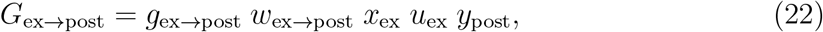

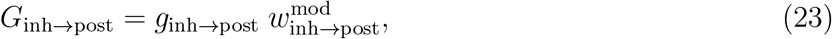

where *g*_pre→post_ is the maximal synaptic conductance per single spine for connections between postsynaptic (‘post’) and presynaptic (‘pre’) neurons; *x, u*, and *y* are short-term plasticity states; *w*_ex→post_ is the excitatory synaptic weight; and 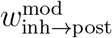 is the state-dependent modulation of the inhibitory weight.

### Addition and removal of spines

Synaptic weights, *w*_pre→post_, represent postsynaptic receptor density across spines, including both protein synthesis-dependent stable (late long-term potentiation; late-LTP) and protein synthesis-independent labile (early long-term potentiation; early-LTP) components, with 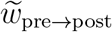 denoting only the stable receptor density (late-LTP) [158]. For excitatory-to-excitatory connections, the variable, *L*_pre→post_, represents the number of spines (each holding up to one unit of receptor density when at peak), capping the maximum allowable potentiation in *w*_pre→post_.

During potentiation attempts in an excitatory-to-excitatory connection, if all spines reach peak receptor density (i.e., *w*_pre→post_ approaches *L*_pre→post_) but are not fully stabilized (i.e., 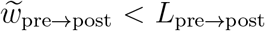), no potentiation occurs. However, if the receptor density is stabilized (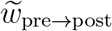 approaches *L*_pre→post_) through protein-dependent processes, a new spine is added by incrementing *L*_pre→post_ by 1, potentially by inducing spine remodeling, allowing further receptor density increases. This ensures receptor density is sequentially allocated across spines by saturating them to full capacity (1 unit each) before distributing any remainder to an additional spine.

Conversely, during depression, as *w*_pre→post_ decreases, the receptor density in one or more spines may reach zero. Upon stabilization of these changes (reflected in 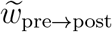), the number of spines is adjusted according to

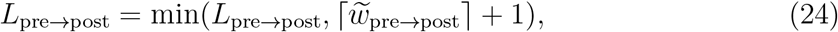

where the ceiling function, 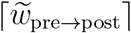, determines the minimum number of spines required to accommodate the stable receptor density. Adding +1 permits at most one spine with zero stable receptor density, effectively deleting excesses in *L*_pre→post_ that would yield more than one such spine. The min operation prevents premature addition of new spines; for instance, if *L*_pre→post_ = 1 and 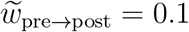, *L*_pre→post_ remains 1, avoiding interference with the addition mechanism. This mechanism facilitates the removal of redundant non-functioning spines while preserving the minimal viable synaptic structure and mirrors empirical observations in biological synapses where repetitive long-term depression leads to pruning of spines with stably reduced receptor density [85].

### Excitatory-to-excitatory long-term plasticity

Synaptic weights, *w*_pre→post_, are updated using a triplet spike-timing-dependent plasticity rule [78]. Upon a presynaptic spike, weights to postsynaptic neurons are depressed proportionally to the product of the fast postsynaptic trace, 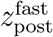, and a term combining pair-based and triplet-based depression factors:

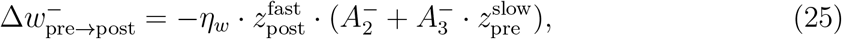

where 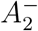 is the pair-based depression amplitude, and 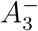 is the triplet-based depression amplitude.

Conversely, upon a postsynaptic spike, weights from presynaptic neurons are potentiated similarly:

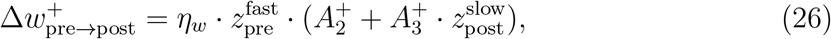

where 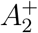 is the pair-based potentiation amplitude, and 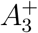 is the triplet-based potentiation amplitude.

Synaptic traces are modeled as low-pass filters of spike trains [158], evolving according to:

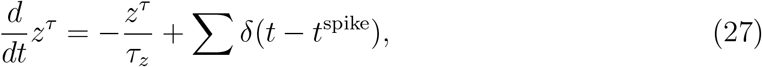

where *z*^*τ*^ is the trace with time constant *τ*_*z*_ (in ms) (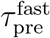 and 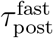 for fast traces capturing recent activity, 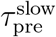 and 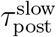 for slow traces capturing earlier activity). In the update rules, the fast presynaptic trace, 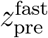, is evaluated at the current spike time *t*, while the other three traces 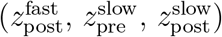 are evaluated just prior to *t* to exclude the current spike. This offset ensures that depression reflects only presynaptic spikes preceded by postsynaptic activity, potentiation dominates without cancellation when spikes coincide, and slow traces incorporate solely prior spikes.

The learning rate, *η*_*w*_, is scaled such that plasticity is enhanced only when both presynaptic and postsynaptic firing rates—*f*_pre_ and *f*_post_ (computed as synaptic traces with a time-constant of 1 s; *z*^1^)—significantly exceed their baseline rates [220, 221], *f*_ref,pre_ and *f*_ref,post_, by a factor of *c*_*η*_:

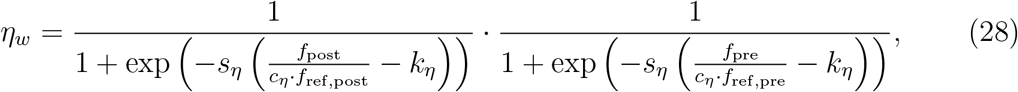

where *s*_*η*_ is the steepness parameter, and *k*_*η*_ is the midpoint threshold.

### Inhibitory long-term plasticity mechanisms

#### Dynamic feedforward inhibition

To preserve excitation-inhibition balance, modifications to excitatory-to-excitatory synaptic weights are mirrored by coordinated adjustments in the feedforward inhibitory circuitry [79]. When the weight, *w*_pre→post_, changes by 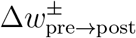 (positive for potentiation, negative for depression), the excitatory-to-inhibitory weights, 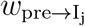, and inhibitory-to-excitatory weights, 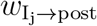, involving the *M* inhibitory interneurons that form feedforward paths are updated according to:

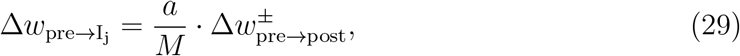

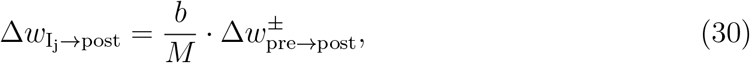

where *j* = 1 to *M* indexes the interneurons, and *a* and *b* are scaling constants. These coupled updates ensure proportional scaling of inhibitory drive in response to excitatory changes.

#### Dynamic feedback and lateral inhibition

In non-background regions (entorhinal, hippocampal, and cortical), we employ distinct inhibitory neurons to mediate feedback [222] and lateral inhibition [223]. Each cell assembly in these regions is assigned one assembly-specific inhibitory neuron (ASIN), which provides feedback inhibition to excitatory neurons within its assembly and lateral inhibition to excitatory neurons in other assemblies, promoting winner-takes-all dynamics [80, 224]. The weights from ‘post’ to its ASIN, *w*_post→ASIN_; from the ASIN to each neuron in its assembly, *w*_ASIN→self_, for feedback inhibition; and from the ASIN to each neuron in other assemblies, *w*_ASIN→other_, for lateral inhibition are updated according to:

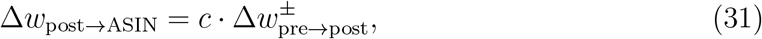

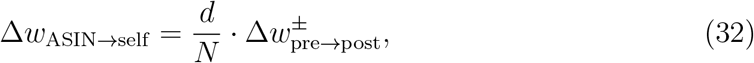

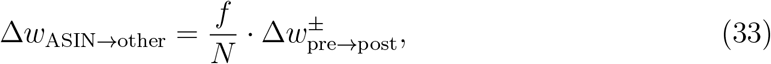

where *N* is the number of neurons in the assembly to which ‘post’ belongs; *c, d*, and *f* are scaling constants.

#### K^+^-dependent disinhibition modulating factor

During sleep, extracellular potassium concentration ([K^+^]_o_) decreases [86], contributing to reduced neuronal excitability through membrane hyperpolarization [225]. Concurrently, acetylcholine levels drop to minimal values during deep sleep [57], unblocking M-current in interneurons and reducing their responsiveness to synaptic inputs [50]. In parallel, norepinephrine levels, reduced during sleep [226], diminish GABAergic inhibition [56], whereas high norepinephrine during wakefulness enhances it via adrenergic receptors [227].

To model this sleep-modulated weakening of inhibitory synaptic transmission without structural plasticity, we introduce a multiplicative modulating factor, denoted as *f*_*I*_([K^+^]_o_), evolving as a sigmoid function to ensure a smooth transition based on [K^+^]_o_ levels:

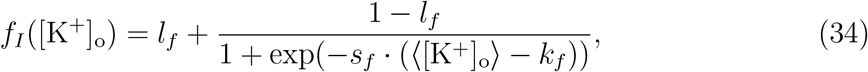

where ⟨[K^+^]_o_⟩ is the average [K^+^]_o_ in the background region, *l*_*f*_ is the minimum scaling value reached during slow-wave sleep (varying by region: MEC, DG, CA3, CA1, CR, Background), *s*_*f*_ (in mM^−1^) is the steepness parameter of the sigmoid, and *k*_*f*_ (in mM) is the midpoint potassium concentration where the factor is halfway between *l*_*f*_ and 1. This factor scales the baseline I→E and I→I synaptic weights:

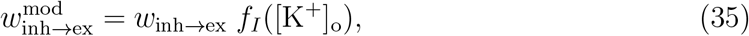

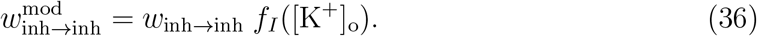

### Short-term plasticity

Short-term synaptic plasticity in excitatory synapses is modeled using the Tsodyks-Markram framework for the presynaptic dynamics [81, 158],

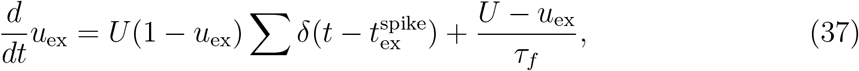

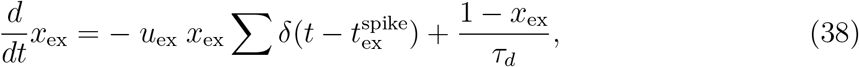

where *u*_ex_ is the neurotransmitter release probability, enhanced upon each presynaptic excitatory spike, relaxing to baseline *U* with time constant *τ*_*f*_ (in ms); *x*_ex_ is the available synaptic resources after neurotransmitter release, depleted by vesicle pool reduction, recovering with time constant *τ*_*d*_; and *δ* denotes Dirac delta functions at spike times.

To prevent runaway excitation during high postsynaptic firing, we model calcium-dependent spike-frequency adaptation at the postsynaptic end, inspired by mechanisms described in the literature [228, 229]. High postsynaptic spiking elevates intracellular calcium via action potential-triggered activation of voltage-gated calcium channels, leading to adaptation that reduces synaptic responsiveness [228]. In our model, this process attenuates the responsiveness of postsynaptic excitatory and inhibitory neurons through the following dynamics:

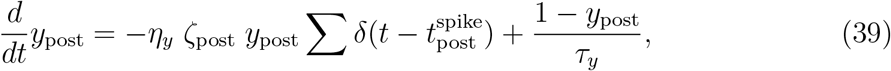

where *y*_post_ is the postsynaptic adaptation factor, depleted in an activity-dependent manner during spiking and recovering to 1 with time constant *τ*_*y*_; *η*_*y*_ (in s^−1^) is the baseline depletion rate; 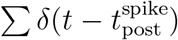 triggers updates at spike times; and *ζ*_post_ is the activity-dependent scaling factor, defined as

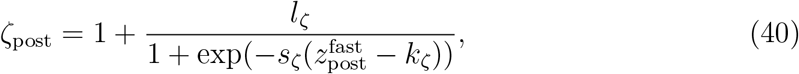

with amplitude *l*_*ζ*_, slope *s*_*ζ*_, and midpoint threshold *k*_*ζ*_.

### Protein-mediated synaptic stabilization

To capture the stabilization of labile synaptic changes into persistent forms, reflecting synaptic tagging and capture mechanisms [230, 231], we model protein-mediated stabilization of synaptic changes. Long-term synaptic plasticity, such as potentiation (LTP) or depression (LTD), involves early phases (early-LTP or early-LTD) with rapid, labile alterations in receptor density (e.g., AMPA receptor trafficking), but these decay without protein synthesis [84], governed by:

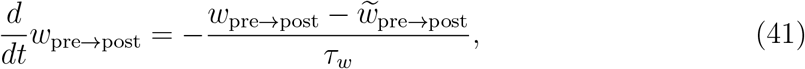

where the total synaptic weight, *w*_pre→post_, decays toward the stable component 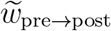 with *τ*_*w*_ as the decay time constant for labile changes.

Activity-dependent intracellular calcium triggers gene expression and synthesis of plasticity-related proteins (PRPs) through pathways like CaMKK/CaMKIV-CREB [232], which are consumed during stabilization. Thus, we model PRP dynamics as:

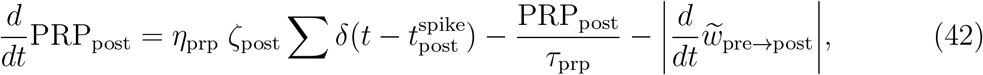

where *η*_prp_ is the baseline PRP increment, *τ*_prp_ is the PRP decay time constant, and 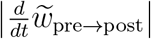 accounts for PRP consumption proportional to stabilization.

These PRPs are captured by tagged synapses to stabilize labile synaptic changes into late-LTP or late-LTD [230, 233], updating the stable weight, 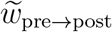, toward *w*_pre→post_ at a rate scaled by available PRP:

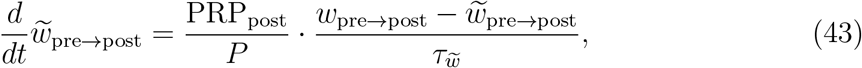

where 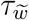 is the stabilization time constant and *P* is the total number of presynaptic neurons.

### Network architecture

We develop an entorhinal-hippocampal-cortical network model using Hodgkin-Huxley-type neurons. The model comprises the medial entorhinal cortex (MEC; 16 excitatory, 7 inhibitory neurons), dentate gyrus (DG; 16 excitatory, 7 inhibitory), CA3 (32 excitatory, 9 inhibitory), CA1 (16 excitatory, 8 inhibitory), cortical regions (CR; 16 excitatory, 8 inhibitory), and a background population representing unmodeled areas (240 excitatory, 60 inhibitory).

In non-background regions (MEC, DG, CA3, CA1, CR), excitatory neurons form four cell assemblies per region. Each assembly pairs with one ASIN that delivers feedback inhibition to its own assembly and lateral inhibition to the other three. This reserves four inhibitory neurons per region; they mutually inhibit with a weight of 1 and lack self-inhibition. Remaining inhibitory neurons process feedforward inhibition.

Background neurons are recurrently connected, with each receiving afferents from approximately 48 randomly selected excitatory and 12 randomly selected inhibitory background neurons. Synaptic weights, *w*_pre→post_, follow a log-normal distribution with parameters 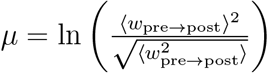 and 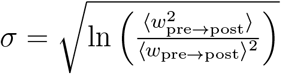 to match specified ensemble averages, where ⟨*w*_pre→post_⟩ is the mean synaptic weight and 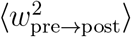 is the mean squared synaptic weight, fixed for each connection type (E→E, E→I, I→E, I→I), generating asynchronous irregular activity with realistic firing rates [88].

Excitatory neurons in non-background regions receive inputs from approximately 48 random excitatory background neurons, with weights drawn from the same log-normal distribution. Inhibitory weights for feedforward, feedback, and lateral inhibition depend on corresponding excitatory-to-excitatory updates and are adjusted as described previously. Two inhibitory neurons per non-background region mediate feedforward inhibition from the background.

Inter-regional projections mirror anatomical connectivity [183]: MEC projects to DG, CA3, and CA1 (one inhibitory neuron per target region for feedforward inhibition); DG to CA3 (one in CA3); CA3 to CA1 (one in CA1); and CA1 to CR (one in CR) [234]. Intra-regional recurrent connections are implemented in MEC [235], CA3 [183], and CR [236–238] (one each in MEC, CA3, and CR). Recurrent connectivity in DG and CA1 is not modeled for simplicity, as it is sparse and predominantly inhibitory in nature [80, 239, 240]. MEC→DG/CA3/CA1, CA3→CA1, and CA1→CR projections are performed assemblywise, where a bundle from a cell assembly projects to a respective bundle of a cell assembly in the receiving area in a one-to-one manner [234, 241, 242]. These projections have initial effective synaptic weights (2 for MEC→DG/CA3/CA1 and 6 for CA3→CA1) [243–245] that assist in signal propagation to downstream regions when MEC is stimulated during the encoding phase. In contrast, recurrent connections in MEC, CA3, and CR are all-to-all without self-connections, while DG→CA3 is all-to-all. These connections—critical for sequential engram encoding—have ∼zero initial weights [246], indicating no pre-stored engrams. Before incorporating CA1→CR projections, parameters *g*_pre→post_ are optimized to yield mean firing rates (⟨*z*^1^⟩) of ∼1 Hz for background excitatory neurons and ∼6 Hz for inhibitory neurons [88], producing asynchronous-irregular dynamics (CV_ISI_ = 1.03, CC = 0.004 [247]; excitatory *z*^1^: mean 1.00 Hz, variance 1.03 Hz^2^; inhibitory *z*^1^: mean 6.00 Hz, variance 13.96 Hz^2^ [88]). Additionally, *w*_inh→ex_ is tuned for entorhinal, hippocampal, and cortical regions to maintain ⟨*z*^1^⟩ of ∼1 Hz. Subsequently, CA1→CR excitatory projections are added with weight of 2 and accompanied by strong feedforward inhibition inserted with *b* equals 0.15 [92–95], resisting functional engram maturation in CR before offline consolidation [38].

In the model, inter-regional delays incorporate theta phase progression to create temporal windows for local circuit computations [82, 248, 249]. These delays are set illustratively below empirical upper limits: 15 ms from medial entorhinal cortex (MEC) to dentate gyrus (DG), 20 ms to CA3, and 25 ms to CA1; 15 ms from DG to CA3 and from CA3 to CA1; and 35 ms from CA1 to cortical regions (CR). Intra-regional inter-assembly delays within MEC, CA3, and CR are assumed to be 20 ms to enable sequential activations during offline replays [103, 189, 250, 251]. Intra-assembly delays are 1 ms. Delays from excitatory-to-inhibitory neurons (for feedback/lateral inhibition), from inhibitory-to-excitatory neurons, and from inhibitory-to-inhibitory neurons are 0 ms.

### Stimulus-induced phase precession

To simulate the encoding of location sequences, we implement stimulus-induced phase precession in medial entorhinal cortex (MEC) neurons relative to 8-Hz cycles whose source is arbitrary. This arbitrary source was chosen to ensure generality across species, irrespective of the specific origin of the cycle to which phase precession is locked: continuous 4–12 Hz hippocampal theta oscillations in the local field potential of rodents [252], ∼8-Hz wing-beat rhythms in bats despite absent hippocampal theta [33], and heterogeneous 2–10 Hz fluctuations in the hippocampus of primates (including humans) during both transient theta bouts and the more prevalent non-rhythmic periods [253, 254]. Four virtual locations (A–D) are traversed in 1.125 s, spanning nine 8-Hz cycles. The 16 MEC excitatory neurons are divided into four cell assemblies (four neurons each), with assemblies 1–4 representing locations A–D, respectively [255, 256].

Stimulation follows a phase-precession pattern across 8-Hz cycles:

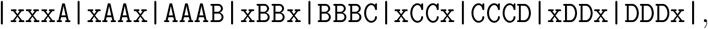

where each segment between vertical bars represents one 8-Hz cycle, ‘x’ denotes no stimulation, and ‘A’–’D’ indicate stimulation of the corresponding assembly. This pattern replicates the empirical shift of location-tuned firing to earlier phases [257]. For repeated explorations of locations A–D, the stimulation pattern is repeated with a gap of one 8-Hz cycle following each sequence, such that four complete explorations occur over 40 8-Hz cycles, totaling exactly 5 s.

Each stimulation event consists of a burst of three current pulses with 6-ms inter-pulse intervals, each designed to elicit one spike. For each pulse, external current is applied for up to 2 ms; if a spike occurs sooner, the current is immediately removed to ensure exactly one spike per pulse.

### Local field potential proxy estimation

We estimate the proxy for local field potentials (LFPs), Φ^*R*^(*t*) (in mV), as proportional to the average post-synaptic current of the excitatory neurons in a region R [258], using a point-source volume conduction approximation [259]:

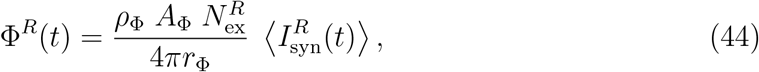

where 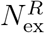 is the number of excitatory neurons in R (plus background neurons if R≠ Background), with *R* ∈ {MEC, DG, CA1, CA3, CR, Background}, and 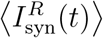 is their average post-synaptic current density at time 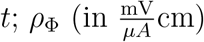 is the extracellular resistivity [260, 261], *r*_Φ_ (in cm) is the typical electrode distance [259, 262], and *A*_Φ_ (in cm^2^) is the effective membrane area per neuron, incorporating dendritic contributions based on standard pyramidal cell morphologies [263, 264]. These parameters yield empirically consistent LFP amplitudes [265, 266].

### Measures of spike train variability and neuronal synchrony

We quantify the irregularity in neuronal firing patterns using the inter-spike interval coefficient of variation (CV_ISI_), calculated as

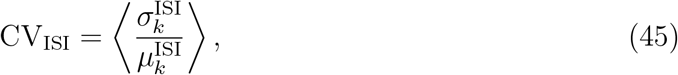

where the angle brackets represent averaging across the entire neuronal population, 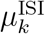 denotes the average inter-spike interval for the *k*-th neuron, and 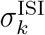 is its corresponding standard deviation.

To assess the degree of coordinated activity among neurons, we employ the mean pairwise correlation coefficient (CC), defined following [87] by

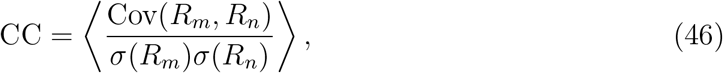

with the averaging performed over every unique pair of neurons. Here, *R*_*m*_ indicates the spike count for neuron *m* within non-overlapping time bins of 5 ms, Cov(*R*_*m*_, *R*_*n*_) is the covariance between the counts for neurons *m* and *n*, and *σ*(*R*_*m*_) represents the standard deviation of the spike counts for neuron *m*.

### Detection and extraction of sequential cell assembly replays

For each of the four CA1 assemblies, we compute the average synaptic trace across members using a time constant *τ* = 25 ms (⟨*z*^0.025^⟩). Peaks in this trace, corresponding to the maxima of smoothed activation events, are detected to timestamp each assembly activation. To extract sequential replays in the order 1 → 2 → 3 → 4 (including partial chains of length ≥ 1), we link each peak in assembly *k* to the nearest subsequent peak in *k* + 1 within 125 ms. Longest chains (up to length 4) are constructed first, followed by shorter chains from remaining peaks, with no reuse to avoid overlaps or redundant counts.

### Estimation of mean ripple power

Ripple-associated replay power, *P*_*e*_ (in mV^2^), in the simulated CA1 LFP time series, Φ, is estimated per replay event as the mean-squared amplitude:

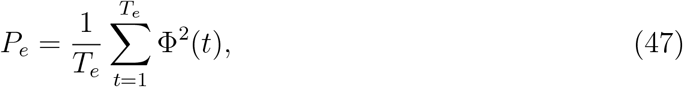

where *T*_*e*_ is the number of time points in the replay event. The mean power is then obtained by averaging *P*_*e*_ across replay events.

### Numerical Integration

The differential equations were discretized by approximating *dt* as Δ*t* = 0.025 ms and *dW* as Δ*W* ∼ 𝒩 (0, Δ*t*). Eq. (1), incorporating stochastic noise, was solved using the stochastic Heun method [267], which provides second-order weak (distributional) convergence and first-order strong (pathwise) convergence. Eqs. (5–7,14–16,20,21) were solved using the deterministic Heun method [268], which provides second-order convergence. Eqs. (27,37– 39,41–43) were solved using the forward Euler method [269], which provides first-order convergence.

The values and descriptions of all parameters are provided in Table 1.

**Table 1.**
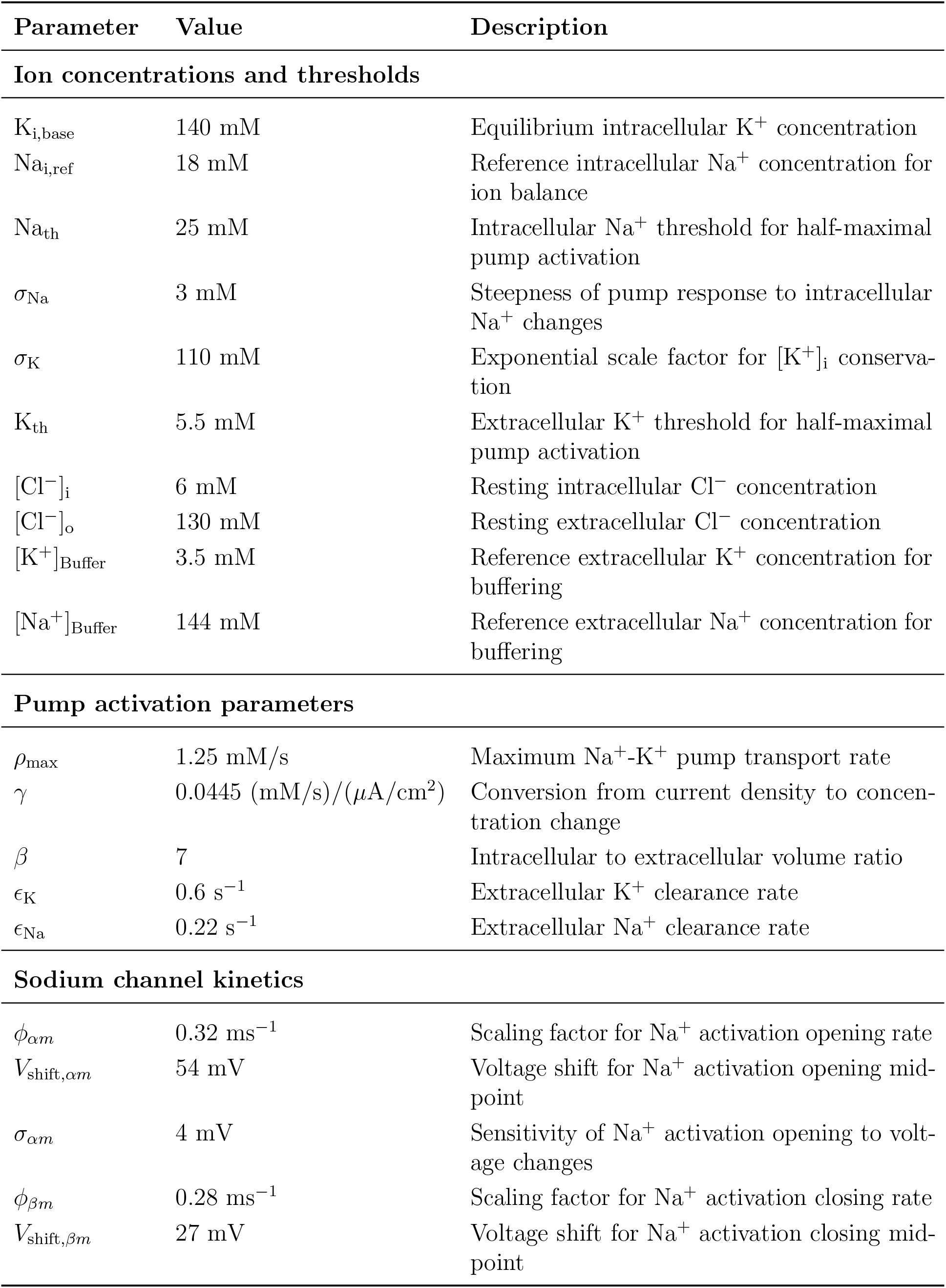
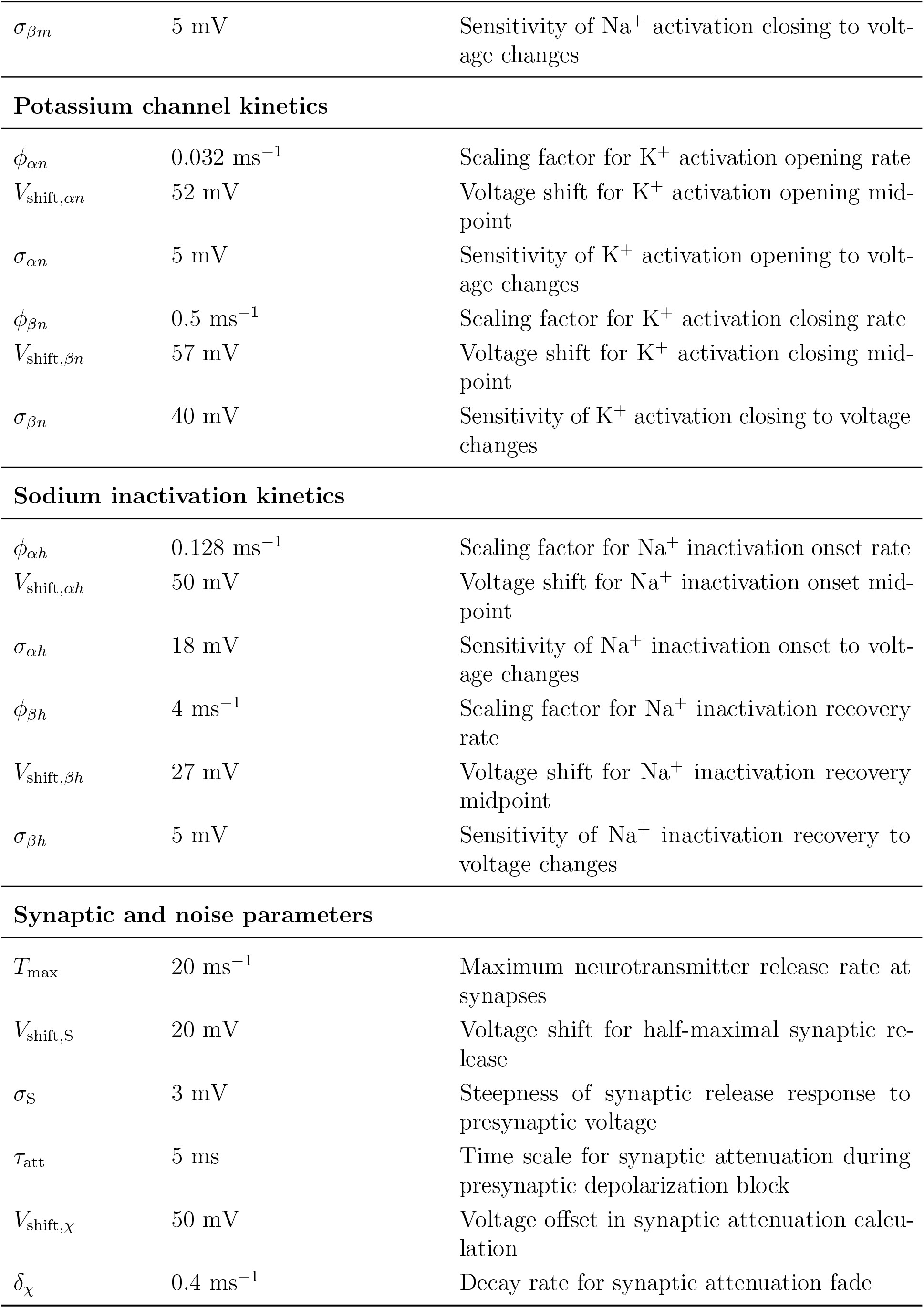
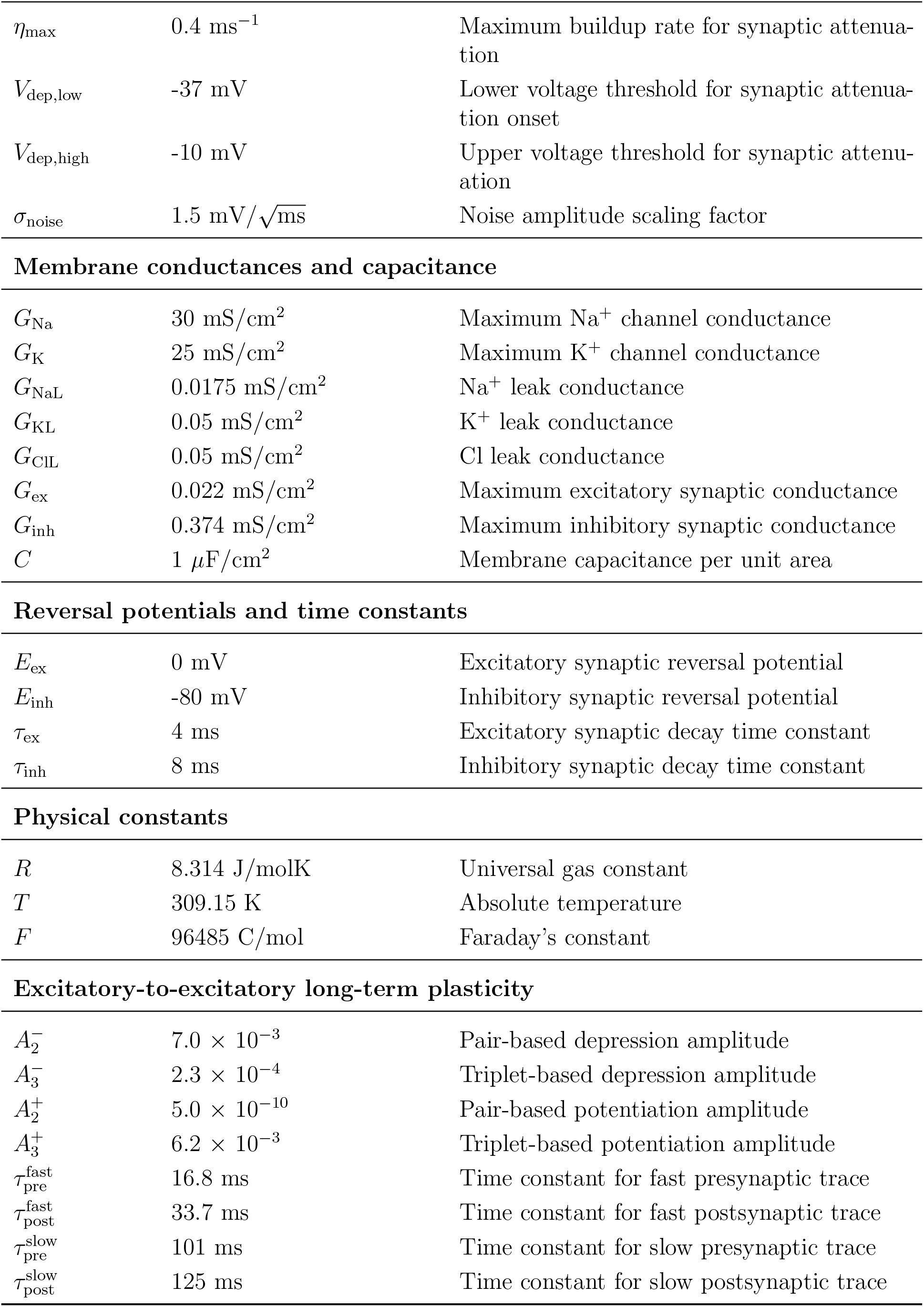
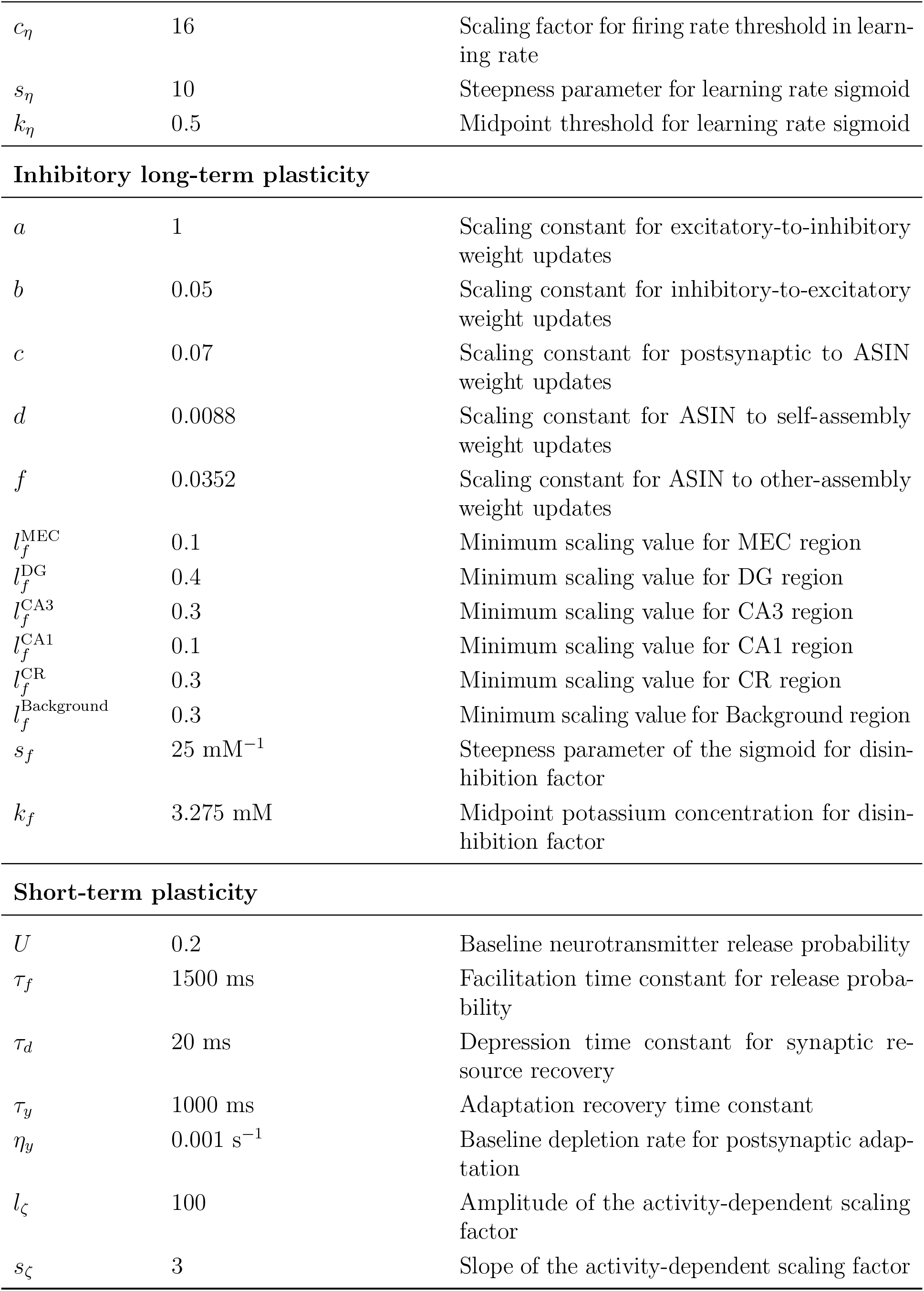
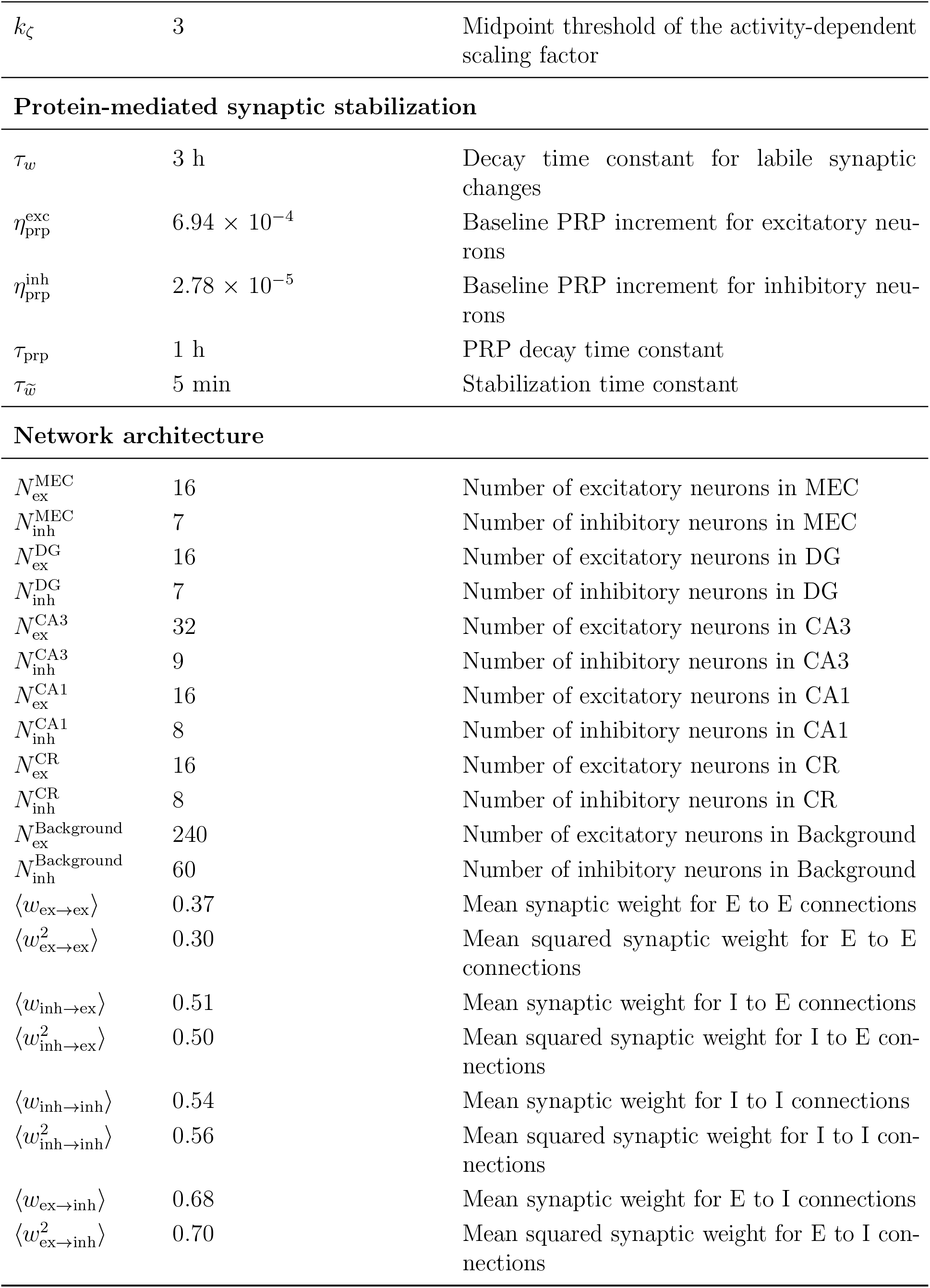
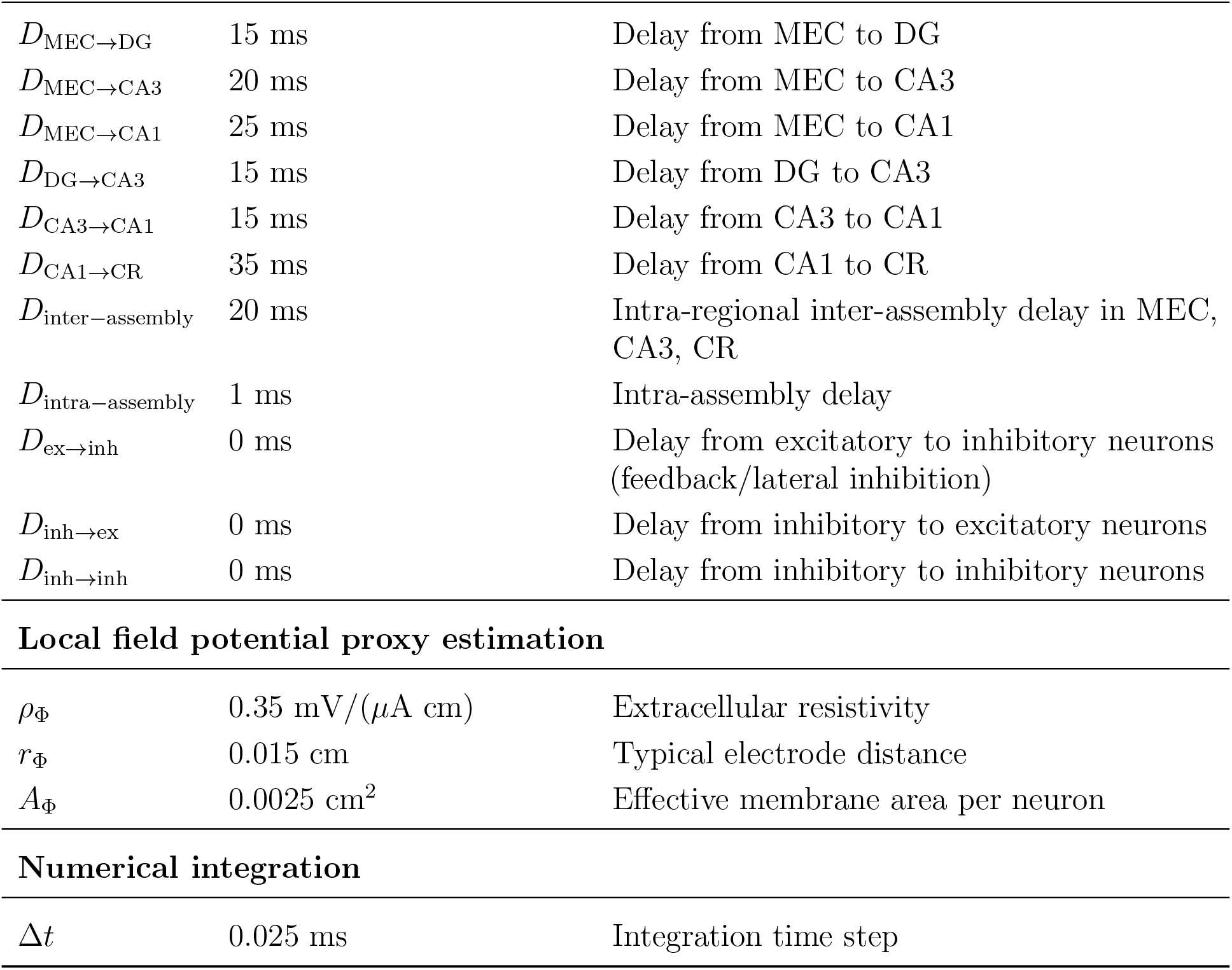
Parameter values and descriptions.

## Acknowledgments

I am grateful to my immediate and extended family for their selfless, invaluable support.

